# Non-target effects on the soil microbiome of a plant root exudate biocontrol for Fusarium wilt

**DOI:** 10.1101/2025.02.10.637461

**Authors:** Eliana Torres-Bedoya, David J. Studholme, Rachel Warmington, Daniel P. Bebber

## Abstract

Crop rotation and intercropping with *Allium* spp. (e.g. garlic, onion) are recognized as biological controls of Fusarium wilt in several crops. However, the non-target effects of this strategy on the soil microbiome are largely unknown. We evaluated the effect of cultivating *Musa basjoo*, *Allium tuberosum* (Chinese leek), and their co-cultivation on the rhizosphere and bulk soil microbiome under glasshouse conditions. We do not report the impact of Chinese leek on Foc abundances, as this has been previously reported. The bacterial and fungal communities in the rhizosphere and bulk soil varied depending on the plant type. The microbiomes of allium and bananas were consistently dissimilar, and the co-cultivation treatment contained constituent taxa from both. However, the effect of allium on soil community composition was outweighed by that of bananas in the co-cultivation scenario. Discriminative taxa for allium included members of the genera *Sphingomonas, Microbacterium, Flavihumibacter, Brevundimonas, Pseudolabrys, Ramlibacter, Trichoderma, Mortierella* and *Fusarium*. In the case of bananas, predominant biomarkers encompassed members of *Rhizobacter, Noviherbaspirillum, Pseudarthrobacter, Aquabacterium, Pseudomonas, Tausonia* and *Humicola* Biomarkers and predictions of functional gene abundances suggest that shifts in the soil microbiome induced by allium are correlated with increases in microorganisms exhibiting potential disease suppression and antibiotic or antifungal traits, whereas those in bananas are associated with plant-growth-promoting microorganisms. The biocontrol efficacy of allium co-cultivation may there involve shifts in the soil microbiome as well as direct impacts of root exudate chemistry on *Fusarium* plant pathogens.

## Introduction

The ascomycete fungus *Fusarium oxysporum* (Fo) is the causal agent of a group of soil-borne diseases known as Fusarium wilts (FW), which threaten production of several crops (Y. Gao et al., 2023; Jiskani et al., 2021; Nishioka et al., 2019; Tang et al., 2023; Zhu et al., 2023). Currently, Cavendish banana (*Musa* spp.) cultivation systems are being severely affected by Fo f. sp. *cubense* Tropical Race 4 (TR4), which causes FW of bananas (FWB) (Z. Li et al., 2020; Shafi et al., 2023). A significant correlation between intensive agricultural practice of banana monocropping, FWB and yield depletion was confirmed by Shen et al. in 2018 (Shen et al., 2018).

Crop rotation (culturing different crops on a field every growing cycle and season) and intercropping (growing two or more crops in the same field side by side in the same or overlapping growing season) are two of the oldest and most fundamental agricultural practices (Farina et al., 2018; Z. Li et al., 2020). Crop rotation and intercropping can contribute to soil-borne disease management through diversification of soil microbiomes, changes in crop microclimates, allelopathic effects, and reduction in exposed soil areas, which directly reduce pathogen abundance or favour the emergence of a protective microbial population (De Corato et al., 2020; Hong et al., 2020; Y. H. Huang et al., 2012; Huss et al., 2022; Nishioka et al., 2019; J. Yang et al., 2023). In southeast Asia, crop rotation or intercropping with allium plants (*Allium* spp.) have shown reductions in incidence and severity of diseases such as FW of calabash (Fo f.sp. *lagenariae*), cucumber (Fo f. sp. *cucumerinum*), spinach (Fo f. sp. *spinaciae),* tomato (Fo f.sp. *lycopersici*), and banana (Fo f. sp. *cubense*) (Igarashi et al., 2017; Z. Li et al., 2020; S. Liu et al., 2013; Nishioka et al., 2019; Tang et al., 2023; Yan et al., 2022).

Two hypotheses have been proposed to explain the effect of alliums on FW incidence: allelopathy due to allium bioactive compounds (Y. H. Huang et al., 2012; S. Liu et al., 2013; Nishioka et al., 2019; Wibowo et al., 2015; H. Zhang et al., 2013); and changes in microbiome diversity and soil structure (Igarashi et al., 2017; Nishioka et al., 2022; Wibowo et al., 2015). It has been proposed that antifungal molecules, including volatiles, released by plant roots accumulate in soil pores where they inhibit spore germination and hyphal growth (H. Zhang et al., 2013). However, the strength of evidence to support that hypothesis has been questioned (Nishioka et al., 2019). Changes in soil microbiomes have been associated with disease reductions under intercropping/rotation systems with alliums (Igarashi et al., 2017; Nishioka et al., 2022; Wibowo et al., 2015). Compounds implicated in bacterial-mediated soil suppressiveness have been identified, as well as potential Fo antagonists (Igarashi et al., 2017; Nishioka et al., 2022; Wibowo et al., 2015). Nevertheless, the roles played by the microorganisms implied in those shifts and their relationships with FW incidence remain unclear (Nishioka et al., 2019).

The effect of the rotation system of Chinese leek-banana to manage FWB was first identified in China in 1997, where a reduction of FWB incidence was observed in a field in which Chinese leeks had grown in preceding years (Y. H. Huang et al., 2012). Subsequent investigation demonstrated the inhibitory effect of different organic volatile compounds produced by *A. tuberosum*, such as 2-methyl-2-pentenal, dimethyl trisulfide, dimethyl disulphide, dipropyl disulphide, dipropyl trisulfide, and 2-methoxy-4-vinylphenol against TR4 (Z. Li et al., 2020; H. Zhang et al., 2013; Zuo et al., 2015) The toxic effect of these compounds appears associated with induced cell death by oxidative burst, mitochondrial impairment, and plasma membrane depolarization; however, assays on spores grown on allium extracts have suggested that the inhibitory effect is fungistatic rather than fungicidal (Z. Li et al., 2020; Wibowo et al., 2015; H. Zhang et al., 2013; Zuo et al., 2015). Moreover, the toxic effect of the metabolites released by allium on non-target microorganisms has not been evaluated, and reduced microbial activity has been detected in the rhizosphere of intercropped plants (Wibowo et al., 2015).In this study, we evaluate the effects of the Chinese leek – banana co-cultivation system on bulk and rhizosphere soil microbiomes under glasshouse conditions using soil from the Eden Project, Cornwall, UK. The Eden Project is a large enclosed botanic garden, containing soils created from recycled mineral wastes and composted green and bark wastes. We test the hypotheses that allium reduces the relative abundance of plant pathogenic taxa and increases the relative abundance of taxa associated with biological control of plant diseases. In addition, we determine whether relative abundances of banana core microbiome taxa in bulk soil and the rhizosphere are influenced by the presence of banana and allium plants. It is important to highlight that we did not measure the effect of Chinese leek on the abundance of Foc in the soil, as this has been previously reported in other studies and was not the objective of our study.

## Materials and Methods

### Pot trial

The experiment was conducted using soil from the Eden Project. The Eden Project (Cornwall, UK), owned by the Eden Trust, is an educational charity and social enterprise that encourages the growth of environmental awareness. Two large biomes are represented there: the Rainforest Biome and the Mediterranean Biome. TR4 was detected there in 2009 (https://www.promusa.org/blogpost580-TR4-present-in-the-UK)

Four treatments were established: soil with *A. tuberosum* plants (A), with *Musa basjoo* plants (B), with a co-culture of one banana and two allium plants (AB), and a control without plants (C). Each treatment was replicated ten times in a completely randomized design. *M. basjoo* plug plants, measuring 30 mm in height, and seven-week-old *A. tuberosum* plants grown from seeds were used in the experiment. The trial was carried out in an unheated glasshouse with automatic venting. Temperatures ranged between 15.0 °C and 24.8 °C (mean = 19.4 °C, SD = 2.1 °C), and plants were hand watered as required. The experiment was conducted for twelve weeks; afterwards, the plants were carefully removed from the pots. The bulk soil (Sb) of each pot was homogenized and labelled as bulk, two samples of the soils without plants were included. Soil firmly adhered to the roots was collected and homogenized before sampling and labelled as rhizosphere soil (Sr). The resultant samples (80) were freeze-dried for 26 hours and stored at -20 °C until the DNA extraction was performed.

### DNA extraction

Soil samples were defrosted and sieved (1 mm) to remove large particles of organic matter. DNA extraction was performed using DNAeasy PowerSoil Kit® (Qiagen, UK) according to the manufacturer’s instructions. The integrity and purity of DNA were determined using agarose gel electrophoresis and a spectrophotometer (NanoDrop® One UV-Vis spectrophotometer, Thermo Scientific, UK). DNA concentration was quantified using the Quantifluor® 1 dsDNA System (Promega, UK) and Glomax® Discover Multimode Microplate Reader (Promega, UK). For PCR amplification, all genomic DNA samples were required to have a concentration greater than 20 ng/l and a purity and quality around 1.8 and 2.0 based on A260/280 and A260/230 ratios respectively. DNA samples were frozen and stored at - 20°C.

### Library preparation and sequencing

Sequencing libraries were constructed according to the Exeter Sequencing Service protocol for Illumina MiSeq-based 16S rRNA gene and ITS region studies (http://sequencing.exeter.ac.uk/). DNA samples were standardised to 1 ng/μl by dilution in 10 mM Tris buffer before the 2-step PCR amplification was carried out. The prokaryotic hypervariable regions V1-V3 from 16S rRNA gene were amplified using the primers set 27F (5′-AGAGTTTGATCCTGGCTCAG-3′) and HDA2 (5′- GTATTACCGCGGCTGCTGGCACC-3′). ITS1 region was amplified using the primers ITSF1 (5′- CTTGGTCATTTAGAGGAAGTAA-3′) and ITS2 (5′-GCTGCGTTCTTCATCGATGC-3′); and region ITS2 with ITS3 (5′-GCATCGATGAAGAACGCAGC-3′) and ITS4 (5′-TCCTCCGCTTATTGATATGC-3′).

PCR reactions for each sample were performed in triplicate using 12.5 μl NEBNext high-fidelity PCR master mix (New England Biolabs – M05415/L), 5.0 μl of 1 μM each primer, and 2.5 μl of 1 ng/μl template DNA. Preamplification denaturation was carried out for 3 minutes at 95 °C before 10 cycles of denaturation at 95 °C for 30 seconds, annealing at 55°C (16S rRNA and ITS) for 30 seconds, and initial extension at 72 °C for 30 seconds. The final extension was carried out at 72 °C for 5 minutes and then the amplicons were held at 4 °C. PCR products were cleaned of contaminants and primer dimers using Agencourt AMPure XP beads (Beckman Coulter) following the manufacturer’s instructions. Samples were resuspended in 10 mM Tris buffer.

The second PCR round was conducted for all samples using the Nextera XT In- dex Kit (New England Biolabs, FC-131-1002). Reactions of 25 μl were created using 12.5 μl of NEBNext high-fidelity PCR master mix (New England Biolabs, M05415/L), 2.5 μl of each primer and 7.5 μl of template DNA. Eighteen amplification cycles were carried out in a thermocycler using the previously described conditions. PCR products were cleaned and resuspended as stated above.

Library quantification and quality checks were carried out by the Exeter Sequencing Service (http://sequencing.exeter.ac.uk/). Libraries concentrations were determined using the Quantifluor® 1 dsDNA System (Promega, UK) and Glomax® Discover Multimode Microplate Reader (Promega, UK). Afterwards, the libraries were diluted to 4nM using 10 mM Tris pH 8.5 buffer. The quality of four pooled samples (one per treatment group), was verified using an Agilent 2200 TapeStation with a High Sensitivity D1000 ScreenTape (Agilent Technologies, Germany). 16S rRNA internal positive and negative sequencing controls were included. Sequencing was performed in an Illumina MiSeq next-generation sequencer. Raw sequences were deposited in the Sequence Read Archive database (Sayers et al., 2019) and are accessible via BioProjects (Barrett et al., 2012) PRJNA663296, PRJNA688816 and PRJNA688813.

### Bioinformatic and statistical analysis

Raw FASTQ files were analysed under QIIME 2 (Bolyen et al., 2019) environment (version 2023.2.0). Primer trimming was performed using Cutadapt (Martin, 2011). Sequences were de-noised and merged using DADA2 (Callahan et al., 2016). Samples with less than 1000 reads were excluded from the subsequent analysis. The taxonomic classification was assessed through trained classifiers. For 16S rRNA sequences, an amplicon-region-specific classifier was made using the SSUrRNA SILVA database (Quast et al., 2012) and RESCRIPt (Robeson et al., 2021). The fungal ITS classifier was trained using the UNITE reference database (Nilsson et al., 2019). The commands used are described in the Supplementary file 1.

Ecological community analysis was carried out using the package MicrobiotaProcess (S. Xu et al., 2023) in R. Rarefaction curves were generated to determine whether sequencing depth was sufficient to achieve a reasonable estimate of species richness. The alpha diversity was estimated based on indexes of observed OTU, Chao1, ACE, Shannon, Simpson and Pielou. Principal Coordinate Analysis (PCoA) using the Bray-Curtis dissimilarity matrix was plotted to compare beta diversity differences in microbial communities associated with the treatments. We utilized a permutational multivariate analysis of variance (PERMANOVA, ADONIS) with 9999 permutations to test dissimilarities in the communities’ composition and ascertain their degree of separation. Relative abundances and community compositions at phylum and genus levels were calculated and plotted. Biomarker discovery was performed using alpha values of 0.01 for 16S rRNA and 0.05 for ITS. The functional composition of the 16S rRNA microbiome was predicted using PICRUSt2 (Douglas et al., 2020). The results were analyzed with ggpicrust2 (C. Yang et al., 2023) and LEfSe (Segata et al., 2011). The code used for the complete analysis is described in the Supplementary file 1.

## Results

### Sequence data

After quality filtering and removal of chimeric sequences, a total of 1.6 million bacterial sequence reads were obtained from 79 samples. The length distribution of sequences ranged from 270 to 496 bases (mean = 468 bases). Samples with less than 1000 reads were excluded from the analysis, resulting in a total of 68 samples that clustered into 22050 amplicon sequence variants (ASVs). For ITS1, 5.4 million fungal reads were retained after quality control. Sequences varied from 200 to 346 bases (mean = 263 bases). Following the removal of samples with low reads, 72 samples were retained and clustered into 4829 ASVs. Regarding ITS2, 4.4 million reads of 250 – 416 bases (mean= 349 bases) were preserved for 71 samples. Of those, 61 samples clustered into 6886 ASVs.

Rarefaction analyses showed that the sequencing depth was sufficient to achieve a reasonable estimate of species richness and a good representation of the microbial communities (Figures S1 – S3). A total of 3247 ASVs were identified for 16S rRNA, 3021 for ITS1, and 558 for ITS2.

### Bacterial and fungal community structure

PCoA matrices and ADONIS test indicated Sb and Sr bacterial and fungal communities in A, B, AB, and C were significantly (p < 0.01) dissimilar (Figure 1). In total, the plant type could explain 13.3 – 21.6% of the variation in these microbial communities. The bacterial community structure in Sb showed greater similarity between A and C than with B. Interestingly, the community of AB treatment in Sb and Sr was intermediate between B and A; nevertheless, the effect of A was outweighed by B. Similar results were observed in Sr for ITS1 and ITS2 (Figure 1). In contrast, the fungal community structure in Sb displayed higher similarity between B and C, while A and AB treatments showed higher similarities. Comparable results were obtained with ITS1 and ITS2 regions.

**Figure 1:**
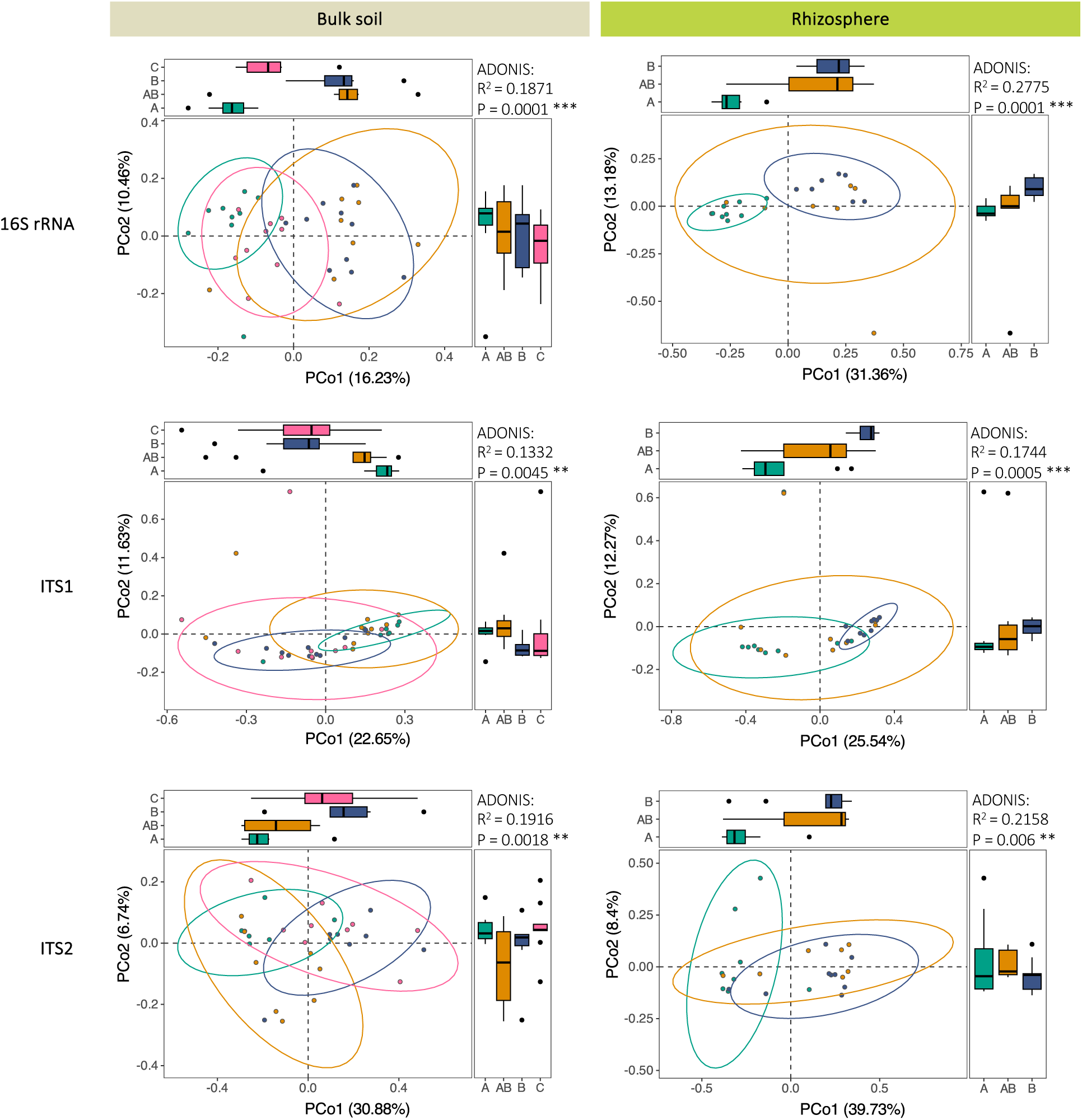
Principal Coordinates Analysis (PCoA) plot and ADONIS test of bacterial (16S rRNA) and fungal (ITS1 and ITS2) community structures bulk soil (Sb) and rhizosphere (Sr) based on the Bray-Curtis dissimilarity matrix. Asterisks indicate significant differences (***: p < 0.001, **: p < 0.01, *: p < 0.05). Allium (A) treatment is colored in green, banana (B) in blue, the co-cultivation allium + banana (AB) in yellow and the control (C) in pink.

### Fusarium oxysporum

Fo was not detected in Sb or Sr using ITS1 region. Thirty ASVs were placed in the *Fusarium* genus without resolution to the species level. Using the ITS2 region, low relative abundances of Fo were found in Sb (C: 0.33 ± 0.67, B: 0.43 ± 0.89, A: 0.65 ± 1.1 and AB: 0.72 ± 1.58) and Sr (AB: 0.58 ± 0.99, B: 0.71 ± 1.35 and A: 1.72 ± 1.21). However, no significant differences (p > 0.05) were found between the treatments (Figure 2).

**Figure 2:**
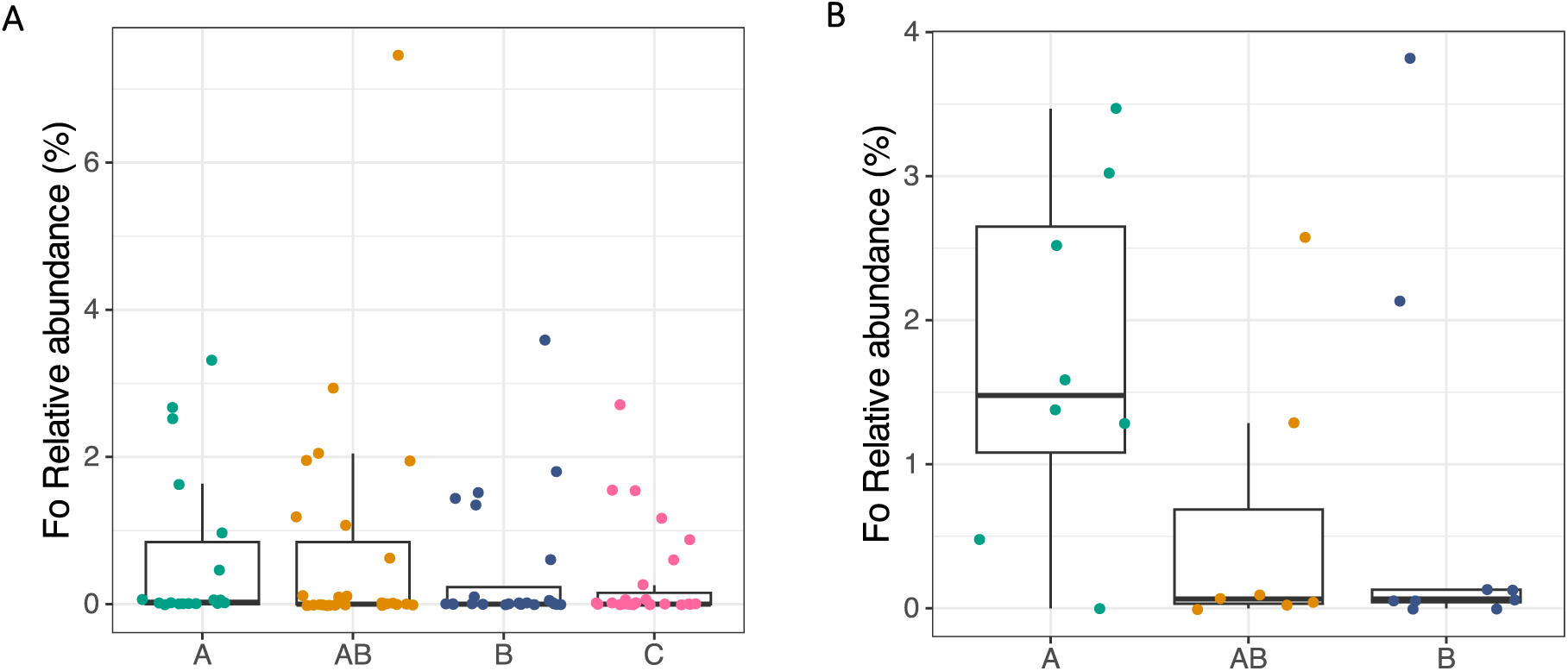
Relative abundance of *Fusarium oxysporum* in bulk soil (Sb) and rhizosphere (Sr) using ITS2 region. Allium (A) treatment is colored in green, banana (B) in blue, the co-cultivation allium + banana (AB) in yellow and the control (C) in pink.

### Community composition

The bacterial community of Sb encompassed 3035 ASVs; 42% were shared between the four treatments (Figure 3). C soil shared 8% of its ASVs exclusively with A, 3% with B and 1% with AB. The AB treatment shared 3 and 4% of its ASVs exclusively with A and B (respectively). The community was dominated by phyla *Proteobacteria* (more abundant), *Actinobacteria*, *Acidobacteriota*, *Chloroflexi*, *Bacterioidota*, *Firmicutes*, *Cyanobacteria* and *Gemmatimonadota* (Figure 3, Table S1). A higher abundance of *Cyanobacteria* (p = 0.012) was found in A. Sr enclosed 2600 ASVs; 21% were shared by all treatments, while 8 and 12% were exclusively shared between A and B, and A and AB, respectively. B and AB shared 17% of their ASVs. The eight more abundant phyla were differentiated from those of Sb only by the presence of *Cyanobacteria*, which in the Sr was less frequent and was replaced by *Patescibacteria* (Figure 4). *Acidobacteriota* was more abundant (p = 0.006) in A. The bacterial genera *Geitlerinema, Nodosilinea* PCC-7104, and *Rhizocola* were significantly (p < 0.05) more abundant in Sb of A, while *Planctopirus* was more abundant in AB, and *Rhizobacter* in Sb of B (Tables S3 and S4). In Sr, Candidatus *Kuenenbacteria*, *Ilumatobacter,* IS-44, RB41, and TRA3-20 were more abundant in A, and *Aquabacterium*, *Asticcacaulis*, and *Corallococcus* in B (Tables S3 and S4).

**Figure 3:**
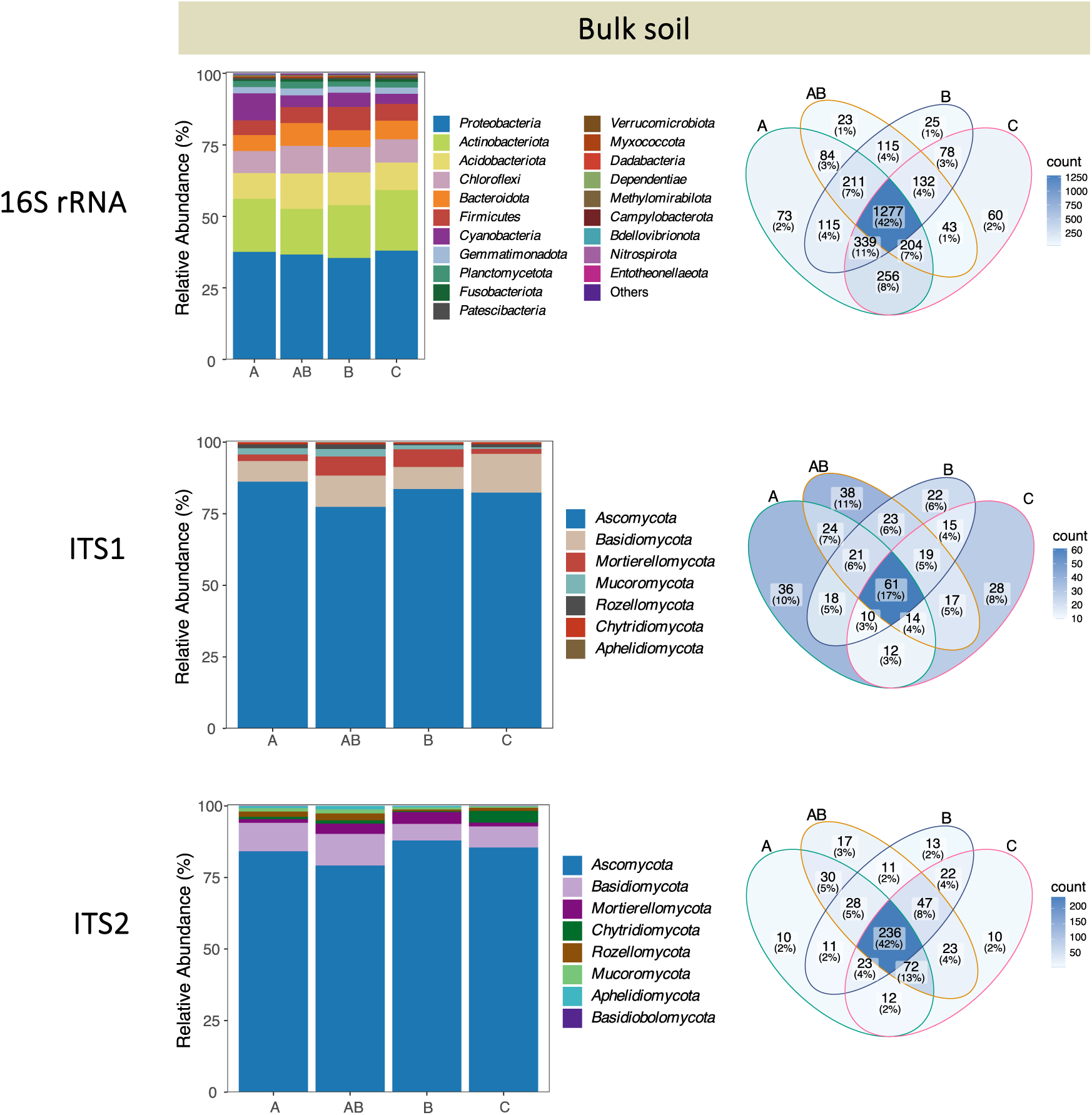
Amplicon sequence variant (ASV) counts and average rarefied relative abundances of the predominant bacterial (16S rRNA) and fungal (ITS1 and ITS2) phyla in bulk soil (Sb) categorized by treatment (allium ‘A’, banana ‘B’, co-cultivation ‘AB’ and control ‘C’).

**Figure 4:**
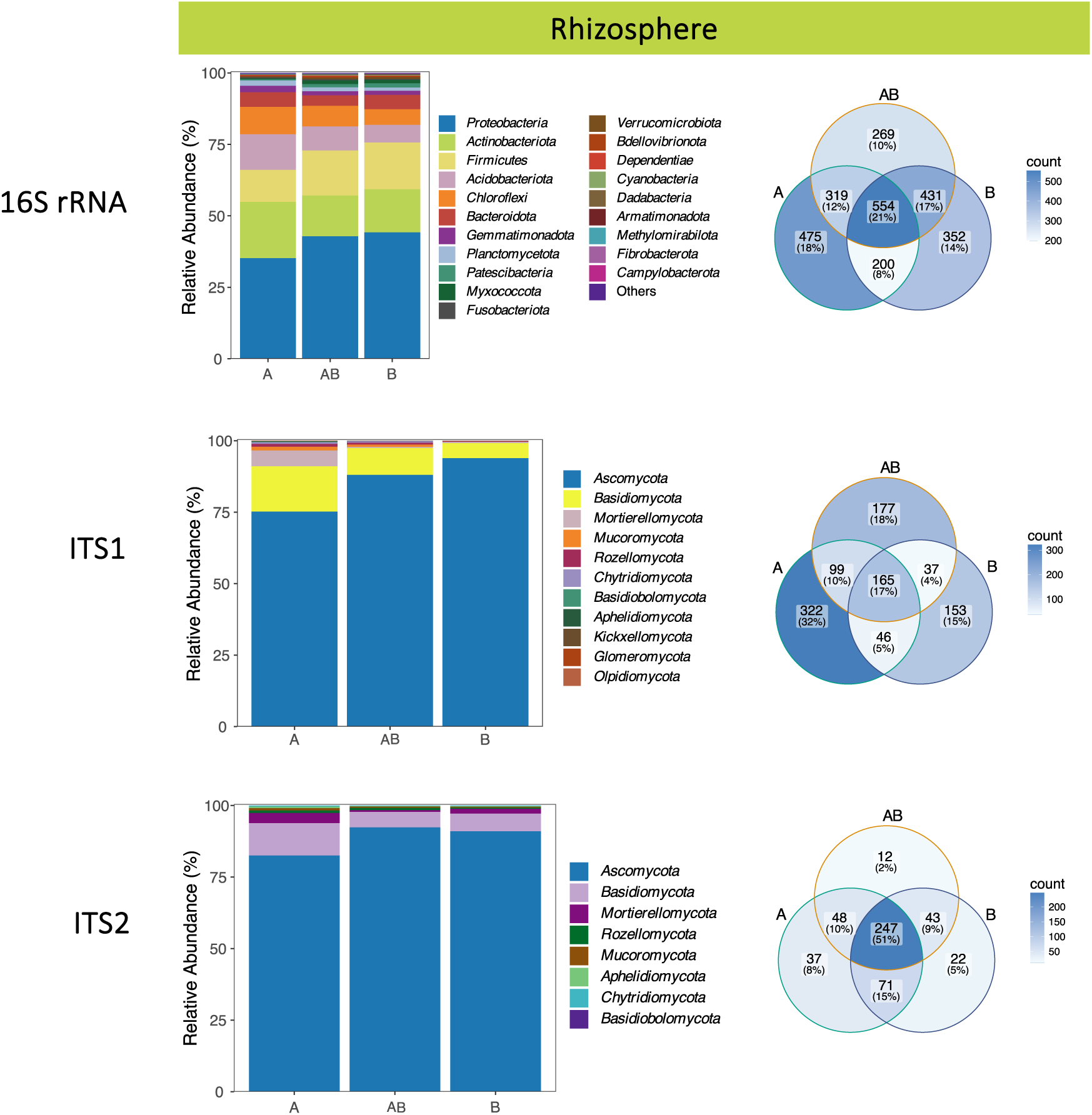
Amplicon sequence variant (ASV) counts and average rarefied relative abundances of the predominant bacterial (16S rRNA) and fungal (ITS1 and ITS2) phyla in rhizosphere (Sr) categorized by treatment (allium ‘A’, banana ‘B’, co-cultivation ‘AB’ and control ‘C’).

The fungal community composition of Sb varied from 470 identified ASVs with ITS1 to 572 ASVs with ITS2. When using the ITS1 region, 17% of the ASVs were shared across all treatments, while 12%, 15%, and 17% were exclusively shared between C and A, B, and AB, respectively. Additionally, 7% of the ASVs were common only between A and AB, and 6% were shared between B and AB. ASVs were classified into seven phyla: *Ascomycota*, *Basidiomycota*, *Mortierellomycota, Mucoromycota*, *Rozellomycota, Chytridiomycota,* and *Aphelidiomycota*; no significant difference (p > 0.05) was found in their relative abundance between the treatments. With ITS2 region, 42% of the ASVs were shared by all the treatments; 2% were present in C and A exclusively, and 4% were shared between C and B, and C and AB treatments. 5% of the ASVs were exclusive for A and AB, and 2% for B and AB (Figure 3). Besides the seven phyla detected with the ITS1 region, members of *Basidiobolomycota* were identified with ITS2. No significant differences in phylum abundances were found among the treatments either.

Differences in the abundance of some fungal genera were detected among the treatments. Using ITS1, *Acremonium* and *Aspergillus* were significantly more abundant in Sb of A, while *Apodus, Ascobolus,* and *Tausonia* were more abundant in Sr of B, and *Leptoxyphium, Mortierella,* and *Trichoderma* in Sr of A (Tables S5 and S6). With ITS2, *Scedosporium* and *Wallrothiella* were found to be more abundant in Sb of A, *Apiotrichum, Aquapteridospora, Arthrobotrys, Chloridium, Coprinopsis, Cylindrocladiella, Fusarium, Hyaloscypha, Lasiodiplodia, Microascus, Mortierella, Mycothermus,* and *Scedosporium* in Sr of A, and *Tausonia* and *Zopfiella* in Sr of B and AB respectively (Tables S5 and S6).

A higher number of ASVs were found in Sr when using ITS1 region than ITS2. 999 ASVs were identified with ITS1; 32% of them were found solely in A, while 17% were present in all the treatments. AB shared 10% of the ASVs with A and 4% with B. The eight phyla associated with Sb samples were also found in Sr, which also included *Kickxellomycota, Glomeromycota* and *Olpidiomycota*. Higher relative abundances of *Aphelidiomycota* (p = 0.006) and *Mortierellomycota* (p= 0.020) were found in A compared with B and AB (Table S2). The analysis performed with ITS2 yielded 480 ASVs; 51% were shared between all treatments, 9 and 10% were shared by AB and B, and AB and A, respectively (Figure 4). The same eight phyla described for Sb with ITS2 were retrieved for the Sr. Phyla *Basidiomycota* (p=0.025) and *Mortierellomycota* (p= 0.021) were more abundant in A (Table S2).

Sixty differentially abundant bacterial ASVs (biomarkers) associated with changes in the microbial communities were identified among all treatments (Tables S7 and S8). Of the twenty-five biomarkers for Sb, the five belonging to C were grouped in the genus *Marmoricola* and the families *Sphingomonadaceae* (*Sphingobium* sp. and *Sphingomonas* sp.) and *Rhizobiaceae*. Fifteen ASVs were retrieved for A, with genera *Geitlerinema, Microbacterium, Flavihumibacter, Brevundimonas,* and two members of the *Rhizobiaceae* family as the top discriminative taxa. The genus *Planctopirus* and families *Nitrosomonadaceae, Anaerolineaceae,* and *Pyrinomonadaceae* encompassed the four markers of AB treatment, while the only selected ASV for B belonged to the genus *Rhizobacter*. Thirty-five biomarkers were retained for the Sr community, twenty-seven for A, and eight for B (Tables S7 and S8). The more abundant ASVs associated with A belonged to the genera *Pseudolabrys, Sphingomonas,* and *Ramlibacter,* families *Gemmatimonadaceae, Vicinamibacteraceae,* and orders *Actinomarinales,* and *Gaiellales*. The markers of B rhizosphere were affiliated with *Noviherbaspirillum* sp., *Pseudarthrobacter* sp., *Aquabacterium* sp., *Pseudomonas umsongensis*, families *Ilumatobacteraceae, Saccharimonadales, Micrococcaceae,* and the order *Vicinamibacterales*.

Fungal biomarkers for Sb were detected only for treatment A (Tables S9 and S10); *Acremonium exuviarum* was identified using both ITS regions, and one member of the family *Nectriaceae* was selected with ITS1. In Sr, nine markers were associated with B using ITS1 region. Those ASVs were identified as *Cephalotrichum stemonitis*, *Plectosphaerella cucumerina*, *Apodus deciduus*, *Cephalotrichum stemonitis*, *Solicoccozyma terricola*, *Subulicystidium brachysporum*, *Tausonia pullulans*, *Subulicystidium brachysporum*, and one member of the order *Aphelidiomycetes*. Seven ASVs were obtained for A, including *Acremonium exuviarum*, *Trichoderma erinaceum*, *Mycothermus thermophilus*, two members of the genus *Mortierella* (*Mortierella* sp. and *M. exigua*), one member of the order *Microascales*, and one belonging to the phylum *Ascomycota*. For ITS2 region, one biomarker was associated with BA (*Serendipita* sp.), two with B (*Tausonia pullulans* and *Humicola nigrescens*) and twenty-eight with A. The top discriminating taxa for A were three members of the genus *Scedosporium* (*Scedosporium prolificans, Scedosporium dehoogii,* and *Scedosporium* sp.), *Mortierella ambigua*, *Apiotrichum dehoogii, Mycothermus thermophilus, Coprinopsis annulopora*, one member of the order *Eurotiales* and interestingly, three members of the genus *Fusarium*, including *F. solani*, a pathogen of several agriculturally relevant crops (Tables S5 and S6).

### Alpha diversity

Higher bacterial richness and lower evenness were observed in Sb of A; however, no significant differences were found in diversity metrics. Furthermore, the treatments did not impact bacterial alpha diversity indexes of Sr. Differences in the fungal indexes varied according to the ITS region used for the analysis (Table S11). Using ITS1, Sb exhibited greater evenness in A than in B and C treatments, and no differences were found in richness and diversity indexes. On the other hand, increased diversity was reported in AB compared to B and C, and higher evenness in A/AB vs C and A against B utilizing ITS2. For Sr, A was richer (Observed index), more diverse, and even than B with ITS1; nevertheless, using ITS2, no differences in richness were observed, and A was more diverse and even than the rest of the treatments.

### Functional predictions

In total, 269 and 270 features from the Kyoto Encyclopedia of Genes and Genomes (KEGG) were selected for Sb and Sr (respectively). To identify pathways that might be over or under-represented within each treatment, comparisons of the predicted KEGG abundances were performed. Differences between treatments in Sb were not significant (p > 0.05); nevertheless, thirty-eight differentially abundant KEGG orthologs were identified (p < 0.010, LDA >2.000) for Sr (Table 1). Feature ko00280 (Valine, leucine, and isoleucine degradation) was the only one selected for B. Interestingly, predictions for A suggested overexpression of four metabolic pathways associated with antibiotic biosynthesis: Streptomycin biosynthesis (ko00521), terpenoid backbone biosynthesis (ko00900), ubiquinone and other terpenoid-quinone biosynthesis (ko00130), and polyketide sugar unit biosynthesis (ko00523). Furthermore, several features associated with carbon fixation, amino acids, and vitamin metabolism were also observed. As these outcomes are purely predictive, a cautious approach should be adopted in their interpretation.

**Table 1:** Differentially abundant KEGG orthologs predicted in bacterial microbiomes from the rhizosphere (Sr) through PICRUSt analysis of 16S rRNA sequencing data. ‘Tx’ denotes the treatment where the feature displayed significantly higher abundance.

| Feature | Tx | LDA score | p-value | Pathway name |
| --- | --- | --- | --- | --- |
| ko03010 | A | 3.32 | 0.003 | Ribosome |
| ko00970 | A | 2.95 | 0.003 | Aminoacyl-tRNA biosynthesis |
| ko00190 | A | 2.94 | 0.003 | Oxidative phosphorylation |
| ko00240 | A | 2.93 | 0.004 | Pyrimidine metabolism |
| ko00720 | A | 2.84 | 0.003 | Carbon fixation pathways in prokaryotes |
| ko00230 | A | 2.79 | 0.004 | Purine metabolism |
| ko00290 | A | 2.75 | 0.003 | Valine, leucine, and isoleucine biosynthesis |
| ko03440 | A | 2.74 | 0.003 | Homologous recombination |
| ko00020 | A | 2.71 | 0.003 | Citrate cycle (TCA cycle) |
| ko03060 | A | 2.71 | 0.003 | Protein export |
| ko00400 | A | 2.71 | 0.003 | Phenylalanine, tyrosine, and tryptophan biosynthesis |
| ko00550 | A | 2.67 | 0.003 | Peptidoglycan biosynthesis |
| ko00520 | A | 2.58 | 0.006 | Amino sugar and nucleotide sugar metabolism |
| ko03030 | A | 2.56 | 0.003 | DNA replication |
| ko03020 | A | 2.55 | 0.004 | RNA polymerase |
| ko00770 | A | 2.53 | 0.003 | Pantothenate and CoA biosynthesis |
| ko03430 | A | 2.53 | 0.004 | Mismatch repair |
| ko00195 | A | 2.51 | 0.003 | Photosynthesis |
| ko00900 | A | 2.50 | 0.003 | Terpenoid backbone biosynthesis |
| ko00250 | A | 2.48 | 0.004 | Alanine, aspartate, and glutamate metabolism |
| ko00710 | A | 2.45 | 0.003 | Carbon fixation in photosynthetic organisms |
| ko00280 | B | 2.42 | 0.010 | Valine, leucine, and isoleucine degradation |
| ko00300 | A | 2.41 | 0.004 | Lysine biosynthesis |
| ko04122 | A | 2.41 | 0.003 | Sulfur relay system |
| ko03018 | A | 2.36 | 0.003 | RNA degradation |
| ko00730 | A | 2.35 | 0.005 | Thiamine metabolism |
| ko00521 | A | 2.35 | 0.003 | Streptomycin biosynthesis |
| ko00450 | A | 2.34 | 0.004 | Selenocompound metabolism |
| ko00670 | A | 2.34 | 0.003 | One carbon pool by folate |
| ko00660 | A | 2.33 | 0.003 | C5-Branched dibasic acid metabolism |
| ko03420 | A | 2.31 | 0.003 | Nucleotide excision repair |
| ko00760 | A | 2.24 | 0.004 | Nicotinate and nicotinamide metabolism |
| ko00564 | A | 2.24 | 0.004 | Glycerophospholipid metabolism |
| ko00790 | A | 2.19 | 0.004 | Folate biosynthesis |
| ko00523 | A | 2.15 | 0.003 | Polyketide sugar unit biosynthesis |
| ko00130 | A | 2.09 | 0.004 | Ubiquinone and other terpenoid-quinone biosynthesis |
| ko00340 | A | 2.04 | 0.005 | Histidine metabolism |
| ko00750 | A | 2.02 | 0.003 | Vitamin B6 metabolism |

## Discussion

In the battle against plant pests and diseases, farmers have employed crop rotations and intercropping for millennia (H. Zhang et al., 2013). Fusarium wilts have become increasingly significant as agricultural diseases, threatening the sustainability of numerous crops (Y. Gao et al., 2023; Jin et al., 2019; Jiskani et al., 2021; S. Liu et al., 2013; Nishioka et al., 2019; Zhu et al., 2023). Intercropping or rotations with rice (Ren et al., 2008), wheat (S. Yang et al., 2023), celery (X. Gao et al., 2020), basil (Raza et al., 2022) and mustard (Jin et al., 2019), among other plants, have been used for managing FW in several crops. In the case of bananas, reduction on pathogen loads and FWB alleviation have been observed in intercropping/rotation systems with allium (Z. Li et al., 2020; Wibowo et al., 2015; H. Zhang et al., 2013), pineapple (J. Yang et al., 2023), pepper (Hong et al., 2020), eggplant (Hong et al., 2023), sugarcane (Fan et al., 2022), wax gourd (Fan et al., 2022), pumpkin (Fan et al., 2022), and green manure (J. Yang et al., 2022). In southern China, *A. tuberosum* (Chinese leek) has been used to manage FWB. However, its application is restricted due to limited knowledge about the underlying mechanisms for disease control and its (non-target) effects on soil microbial communities (Nishioka et al., 2019; Wibowo et al., 2015; H. Zhang et al., 2013).

Our study showed that the cultivation of A, B and AB significantly altered bulk soil and rhizosphere bacterial and fungal community structures. Before further discussion of the results, we emphasize that the soil in the Eden Project biomes was created from recycled mineral wastes and composted green and bark wastes, and will differ from soils in actual plantation conditions. A and B microbiomes were dissimilar in all cases and the effect of A in AB community structure was outweighed by that of B in most cases (except in bulk soil fungal community). This result indicates that under intercropping scenarios, the mechanism of action of allium in suppressing Foc can be directly attributed to the release of antifungal compounds into the rhizosphere, thereby reducing Fusarium growth and/or invasion, and not to changes in the microbiome. We observed that allium cultivation alters the microbial properties of the soil and increases the abundances of microorganisms capable of reducing pathogenic Foc. This seems to be the main mechanism contributing to FWB suppression under crop rotation schemes.

Studies evaluating rotations/intercropping systems for FW reduction, such as pineapple (J. Yang et al., 2023), pepper (Hong et al., 2020, 2023), and eggplant (Hong et al., 2023), have shown a decrease in Fo abundance correlated with a reduction in FW incidence. This suggests that the abundance of Fo could potentially serve as an indicator for Foc. Despite the soil used in this study not being infected with Foc, Fo was detected using the ITS2 region. However, no reductions in Fo were observed in treatments A and AB when compared with B. Similar results have been described when managing FWB with biofertilizers (Hong et al., 2023). It has been suggested that this outcome might be explained by the model of pathogen-helper inhibition proposed by Li *et al*. (M. Li et al., 2022), in which the soil microbiome forms a functional barrier against pathogen invasion. Disease suppression is achieved by reducing root-associated bacteria that facilitate pathogen growth/colonization (indirect effect) rather than lessening the pathogen population (direct effect) (Hong et al., 2023; M. Li et al., 2022). Further studies must be performed to determine the exact mechanism of FWB reduction exhibited by allium.

Allium bacterial community exhibited high abundances of *Cyanobacteria* and *Acidobacteriota* in bulk soil and rizhosphere, respectively. High abundances of these two phyla were previously described in soil bacterial community associated with Chinese leek by Huang in 2018 (Y. Huang, 2018) and Gu *et al*. in 2020 (Gu et al., 2020), who reported *Proteobacteria, Acidobacteria, Bacteroidetes*, *Cyanobacteria*, and *Planctomycetes* as the top abundant phyla in allium soil samples (Gu et al., 2020; Y. Huang, 2018). *Cyanobacteria* are known to produce allelochemicals able to inhibit the growth of several fungi and thus have been considered eco-friendly biocontrol agents (Tiwari & Kaur, 2014). Bioinoculants obtained from *Cyanobacteria* have been used to control FW on chili (Yanti et al., 2020), tomato (Prasanna et al., 2013), and pepper (Abdelaziz et al., 2022); nevertheless, Fan *et al*. (Fan et al., 2022) found a positive correlation between the abundances of this taxonomic group and FWB incidence in rotation systems with pepper, sugarcane, wax gourd and pumpkin (Fan et al., 2022) (Table 2). On the contrary, in the same study, a negative correlation was observed between *Acidobacteria* and disease (Fan et al., 2022). Similar results have been found in healthy vs diseased banana plants (Jamil et al., 2022), FWB naturally suppressive soils (Shen, Ruan, Xue, et al., 2015) and treated with Bio-Organic fertilizer as FWB management strategy (Shen, Ruan, Chao, et al., 2015) (Table 2). Intriguingly, *Acidobacteria* are known to be more abundant in soils with low pH (Lauber et al., 2009); nonetheless, it has been widely accepted that in acidic soils a higher FWB incidence is observed due to a detrimental effect on soil diversity (Segura et al., 2021).

**Table 2:** Previous microbiome studies correlating Fusarium wilt incidence and/or *Fusarium oxysporum* relative abundance with taxa selected as biomarkers for A, AB, and B in this study. Taxa are categorized by phylum ‘P’, genus ‘G ‘and ‘ASV’.

| Taxa | Tx-Soil<br>type | Negative Correlation | Positive Correlation |
| --- | --- | --- | --- |
| Cyanobacteria (P) | A – Sb |  | FWB in rotation systems with pepper, sugarcane, wax gourd, and pumpkin (Fan et al., 2022). |
| Acidobacteriota (P) | A – Sr | FWB in rotation systems with pepper, sugarcane, wax gourd, and pumpkin (Fan et al., 2022).<br>FWB in healthy versus diseased banana plants (Jamil et al., 2022).<br>FWB in suppressive versus conducive soils (Shen, Ruan, Xue, et al., 2015).<br>FWB in Bio-Organic treated soils for Foc management (Shen, Ruan, Chao, et al., 2015).<br>FW of banana, cucumber, watermelon and lily in diseased versus healthy soils (J. Yuan et al., 2020) |  |
| Mortierellomycota (P) | A – Sr | FW of banana, cucumber, watermelon and lily in diseased versus healthy soils (J. Yuan et al., 2020). |  |
| Basidiomycota (P) | A – Sr | FWB in pineapple–banana rotation for Foc management (B. Wang et al., 2015) | FWB in banana monocropping versus intercropping with green manures for Foc management (J. Yang et al., 2022) |
| Mortierella (G / ASV) | A – Sr | (G) FWB in diseased versus disease-free soils (D. Zhou et al., 2019).<br>(G) FWB in chili pepper-banana rotation systems versus monoculture (Hong et al., 2020).<br>(G) FWB in pineapple–banana rotation for Foc management (J. Yang et al., 2023)<br>(G) FW of vanilla in suppressive versus conducive soil (Xiong et al., 2017).<br>(ASV) FW of banana, cucumber, watermelon and lily in diseased versus healthy soils (J. Yuan et al., 2020). |  |
| Aspergillus (G) | A – Sb | FWB in residue amendment as a strategy for Foc management (J. Yang et al., 2023)<br>FWB in green manure intercropping versus banana monoculture (J. Yang et al., 2022).<br>FWB in diseased versus disease-free soils (D. Zhou et al., 2019). |  |
| Trichoderma (G) | A – Sr | FWB in diseased versus disease-free soils (D. Zhou et al., 2019). |  |
| *Fusarium* (G / ASV) | A – Sb<br>A – Sr | (ASV) FW of banana, cucumber, watermelon and lily in diseased versus healthy soils (J. Yuan et al., 2020).<br>(ASV) FW of flax in conducive versus suppressive soils (Siegel-Hertz et al., 2018). | (G) FWB in rotation systems with pepper, sugarcane, wax gourd, and pumpkin (Fan et al., 2022).<br>(G) FWB in Bio-Organic treated soils for Foc management (Shen, Ruan, Chao, et al., 2015).<br>(G) FWB in pineapple–banana rotation for Foc management (J. Yang et al., 2023)<br>(G) FW of vanilla in suppressive versus conducive soil (Xiong et al., 2017).<br>(G) FWB in pineapple–banana rotation for Foc management (B. Wang et al., 2015)<br>(G) FWB in green manure intercropping versus banana monoculture (J. Yang et al., 2022).<br>(G) FWB in diseased versus disease-free soils (D. Zhou et al., 2019).<br>(G) FW of cape gooseberry in conducive versus suppressive soils (Bautista et al., 2023). |
| Rhizobacter (G/ASV) | B – Sb | FW of flax in conducive versus suppressive soils (Siegel-Hertz et al., 2018). |  |
| Pseudolabrys sp. (ASV) | A – Sr | (G) FWB in Bio-Organic treated soils for Foc management (Xue et al., 2015). |  |
| Sphingomonas sp. (ASV) | A – Sr/Sb | (G) FWB in chili pepper-banana rotation systems versus monoculture (Hong et al., 2020). | (G) FWB in Bio-Organic treated soils for Foc management (Xue et al., 2015).<br>(G) FWB in rotation systems with pepper, sugarcane, wax gourd, and pumpkin (Fan et al., 2022). |
| Ramlibacter sp (ASV) | A – Sr | (G) FW of cucumber in soils after reductive disinfestation, combined with Biochar and antagonistic microbial inoculation as a management strategy for Fo (Ali et al., 2023). |  |
| Mycothermus thermophilus (ASV) | A – Sr |  | (ASV) FW of banana, cucumber, watermelon and lily in diseased versus healthy soils (J. Yuan et al., 2020). |
| F. solani (ASV) | A – Sr | (ASV) FWB in soils treated with pineapple residues for Foc management (X. Yuan et al., 2021). |  |

Rhizosphere of A showed greater abundances of *Mortierellomycota* (ITS1 and ITS2), *Aphelidiomycota* (ITS1), and *Basidiomycota* (ITS2). Very little is known about the role of *Mortierellomycota* members as biological control agents; nevertheless, several studies have found them increased in healthy versus Fo-infested soils with bananas, vanilla, watermelon, cucumber, potato, rice, ginseng, among others (Y. Wang et al., 2022; Xiong et al., 2017; J. Yang et al., 2023; J. Yuan et al., 2020) (Table 2). Furthermore, members of this phylum have demonstrated the capacity to antagonize pathogens through antibiotic production and improve soil health due to the degradation of a range of toxic organics (F. Li et al., 2018; Melo et al., 2014). Increases of *Basidiomycota* have been described in the rotation system pineapple– banana for Foc management (B. Wang et al., 2015); nevertheless, variable results have been observed when intercropping with different green manures (J. Yang et al., 2022) (Table 2). Even though, *Aphelidiomycota* has been found as a dominant phylum in soil microbiomes and is significantly affected by intensive cropping (K. Sun et al., 2021); by the time of this review, its relationship with FW, Fo or other soil-borne diseases has not been studied.

C treatment showed high abundances of ASVs of the genera *Sphingomonas, Sphingobium* and *Marmoricola*. This group of microorganisms is known for its association with plant disease suppression (including FW and FWB) and antibiotic production (Hong et al., 2020; Jiang et al., 2018; H. Liu et al., 2018). One ASV also belonging to *Sphingomonas* sp. was selected as a biomarker of A, summed with other individuals belonging to genera, *Microbacterium, Flavihumibacter*, and *Brevundimonas*; all of them are beneficial bacteria described as plant growth promoters or implicated pathogen suppression (Deng et al., 2021; L. Lin et al., 2012; Madhaiyan et al., 2010; Naqqash et al., 2020; J. Wang et al., 2023). Nonetheless, one of the markers found belonged to *Geitlerinema*, a genus found in soil microbiomes of grafted tomatoes infected by *Ralstonia solanacearum* (Navitasari et al., 2020); despite this, its role in the disease has not been described. Very little is known about the role of the genus and families associated with AB markers (*Planctopirus* sp., *Nitrosomonadaceae, Anaerolineaceae*, and *Pyrinomonadaceae*) in plant diseases; even so, *Nitrosomonadaceae* have been associated with soil-reduced nitrogen loss and with higher relative abundance in healthy soils versus infested with *Ralstonia solanacearum* when studying bacterial wilt in sesame (Clark et al., 2021; M. Li et al., 2020; R. Wang et al., 2018). On the contrary, *Pyrinomonadaceae* have been found more abundant in soils inoculated with *Tilletia controversa*, a fungal pathogen of wheat (Jia et al., 2022). The only biomarker for B belonged to *Rhizobacter* spp., a genus of nitrogen fixer plant-growth-promoting endophytes, present in flax FW suppressive soils and widely used in the agricultural management of leguminous plants (Adeleke et al., 2021; Bhardwaj et al., 2014; Siegel-Hertz et al., 2018). It has been shown that *Rhizobacter* spp. can act as a biocontrol in plant species other than legumes (Farooq Khan et al., 2022), suggesting that bananas could have selected specific taxa to compete with soil pathogens, as was previously noted by Hong *et al*. (Hong et al., 2023), who found that banana microbiomes contained high abundances of *Formivibrio, Novosphingobium,* and *Burkholderia* spp. (Hong et al., 2023).

In rizhosphere the more abundant biomarkers of A included *Pseudolabrys* sp., *Sphingomonas* sp., and *Ramlibacter* sp., reported in FWB suppressive soils and with a negative correlation with severity in FW of cucumber and sweet pepper disease (Ali et al., 2023; Hong et al., 2023; Xue et al., 2015; L.-N. Zhang et al., 2019). Furthermore, other ASVs belonged to *Vicinamibacteraceae* and *Gaiellales*, which have reported roles in suppressing disease suppressors, soil anti-toxicity and as antioxidants (J. Chen et al., 2020; M. Lin et al., 2023; D. Xu et al., 2022); nevertheless, members of *Gemmatimonadaceae* and *Actinomarinales* have been reported as negatively correlated with plant growth and higher abundance in pathogen-inoculated soils [(Jia et al., 2022; Luo et al., 2022). Overall, the roles of the ASV taxa associated with B were either less known or controversial. Some members of *Noviherbaspirillum* spp., *Aquabacterium* spp., and *Saccharimonadales* have shown antifungal activity, notwithstanding their abundance is associated with higher and lower incidences of several plant diseases (S. Chen et al., 2020; Jia et al., 2022; H. Li et al., 2019; M. Lin et al., 2023; Shang et al., 2021; Y. Wang et al., 2021). The genus *Pseudomonas* has been described as having suppressive effects on Fusarium wilt however, it also includes several pathogenic members (Kwon et al., 2003; Shen et al., 2022). The specific role of *Pseudomonas umsongensis* in agricultural soils remains unknown. Scarce information is available about the correlation of *Ilumatobacteraceae* and *Micrococcaceae* as families and plant diseases; nevertheless, *Pseudarthrobacter* spp. and *Kocuria* spp., members of *Micrococcaceae*, produce molecules with demonstrated anti-fungal (including Fo) activity and harbour plant growth-promoting traits (N. Liu et al., 2020; Setiawan et al., 2022; Shahid et al., 2021).

*Acremonium exuviarum* (ITS1 and ITS2) and one member of *Nectriaceae* (ITS1) were the only ASVs detected in bulk soil, and both were more abundant in A. The biological significance of *A. exuviarum* in soil health or plant diseases is unclear. Members of the genus *Acremonium* have been described as pathogens in several crops; nevertheless, some isolates are responsible for decreases in FW incidence in bananas, tomatoes, and flax (Abe, 2003; Racedo et al., 2013; Tagne et al., 2002; Y. Yang et al., 2021; Yuelian & Qingfang, 2013). Among the more abundant ASVs for B rhizosphere, beneficial effects were previously described for *Tausonia pullulans* and *Humicola nigrescens*, which are associated with reduced incidence of Verticillium wilt and plant growth promotion (Glushakova et al., 2021; Mestre et al., 2017; Ogundeji et al., 2021). On the other hand, *Plectosphaerella cucumerina* (anamorph *Plectosporum tabacinum*) can survive saprophytically in soil by decomposing plant material or act as a necrotrophic pathogen causing sudden death and blight disease in a variety of crops (Pétriacq et al., 2016). Within the biomarkers of A, *Trichoderma erinaceum* has been linked to reductions in FW incidence of tomato, melon, and watermelon (Aamir et al., 2019; Boughalleb-M’Hamdi et al., 2018). On the contrary, *Mycothermus thermophilus* has been positively related to FW-diseased soils in several crops (J. Yuan et al., 2020). It is well known that high abundances of *Fusarium* spp. and *F. solani* (present in rizosphere of A) have been associated with plant diseases; notwithstanding, several non-pathogenic isolates of these microorganisms have been reported as biocontrol of FW and Verticillium wilt (Kaur et al., 2010; Larkin & Fravel, 2002; Sajeena et al., 2020). Additionally, Yuan and colleagues (X. Yuan et al., 2021) noted its relevance in FWB suppression in soils treated with pineapple residues. They suggested due to a similar profile carbon source utilization, *F. solani* reduces TR4 growth by competing for nutrients (X. Yuan et al., 2021). As significant reductions in FWB have been reported in allium - banana systems, further experiments need to be performed to establish the pathogenicity or not of these fusaria.

Overall, the biomarkers analysis confirmed that roots of allium and banana plants induce shifts in soil microbiomes. In bananas, beneficial shifts seem to be, to a greater extent, associated with plant-growth-promoting microorganisms, whereas, in allium, a greater number of ASVs with potential disease suppression and antibiotic/antifungal traits were retained. These results suggest that changes in microbiomes of allium-banana systems could have a role in reductions of FWB. In both treatments, bacteria, and fungi with unknown roles in soil health or plant disease were selected, aside from microorganisms previously described as phytopathogens; nevertheless, further analyses are required to understand the biological meaning of their presence in the soil under the treatments.

There is considerable controversy in the computational approach to link 16S rRNA gene amplicon data with functional genes derived from bacterial reference genomes; nevertheless, due to the costs of evaluating the bacterial community functionality using experimental approaches, a growing number of studies using this type of analysis in environmental samples have been published in recent years (Breitkreuz et al., 2021; Douglas et al., 2020; S. Sun et al., 2020; J. Zhou et al., 2015). Despite the new software, constant updates, and inclusion of more reference genomes, as a consensus, the results of these analyses must be carefully interpreted (Breitkreuz et al., 2021; S. Sun et al., 2020; J. Zhou et al., 2015). Here we used PICRUSt2 (Douglas et al., 2020) software to investigate functional trends associated with the different treatments in the rhizosphere. Most of the differentially expressed predicted traits were found to be overexpressed under A treatment. Intriguingly, among those, four were related to possible antibiotic biosynthetic pathways. Results that match the potential roles of *Pseudolabrys* sp., *Sphingomonas* sp., *Ramlibacter* sp., *Vicinamibacteraceae* and *Gaiellales*, the taxa encompassing the top biomarkers in A rhizosphere. Features associated with carbon fixation, amino acids, and vitamin metabolism also reported in this analysis, could be owing to plant growth promotion; nevertheless, there is insufficient evidence to obtain a reliable conclusion about them.

In summary, our findings indicate that the co-cultivation of allium and bananas does not induce significant changes in the soil microbiome compared to cultivating bananas alone. Consequently, the reduction in the population of Foc reported in earlier studies may be attributed to the allelopathic effect of root exudates produced by allium plants. Conversely, in crop rotation scenarios, cultivating allium plants on their own induces shifts in soil microbial populations that may play a role in reducing the incidence of FWB when susceptible plants are re-planted.

## Declarations

### Ethics approval and consent to participate

Not applicable

### Consent for publication

Not applicable

### Funding

This work supported by the Department of Biosciences, Global Food Security grant no. BB/N020847/1 and EC Horizon 2020 project ID 727624. E. Torres-Bedoya was supported by a doctoral training award funded by Waitrose Agronomy Group/Primafruit, the Ayudar Foundation, and the University of Exeter.

### Availability of data and materials

Raw sequences were deposited in the Sequence Read Archive database (Sayers et al., 2019) and are accessible via BioProjects (Barrett et al., 2012) PRJNA663296, PRJNA688816 and PRJNA688813.

### Competing interests

The authors declare no competing interests.

### Authors’ contributions

D.B. and R.W. conceived and designed the research. R.W. conducted the experiments. E.T. and D.S. conducted the bioinformatic analyses. E.T. analysed the data and wrote the paper with input from D.B. and D.S.

## Acknowledgements

The authors thank Hannah White, Omotola Odetayo and Ben Temperton for contributions to the work.

## Notes

https://www.ncbi.nlm.nih.gov/bioproject/?term=PRJNA663296

https://www.ncbi.nlm.nih.gov/bioproject/?term=PRJNA688816

https://www.ncbi.nlm.nih.gov/bioproject/?term=PRJNA688813

### Competing Interest Statement

The authors have declared no competing interest.

