## Supplementary Material for "Non-target effects on the soil microbiome of a plant root exudate biocontrol for Fusarium wilt"

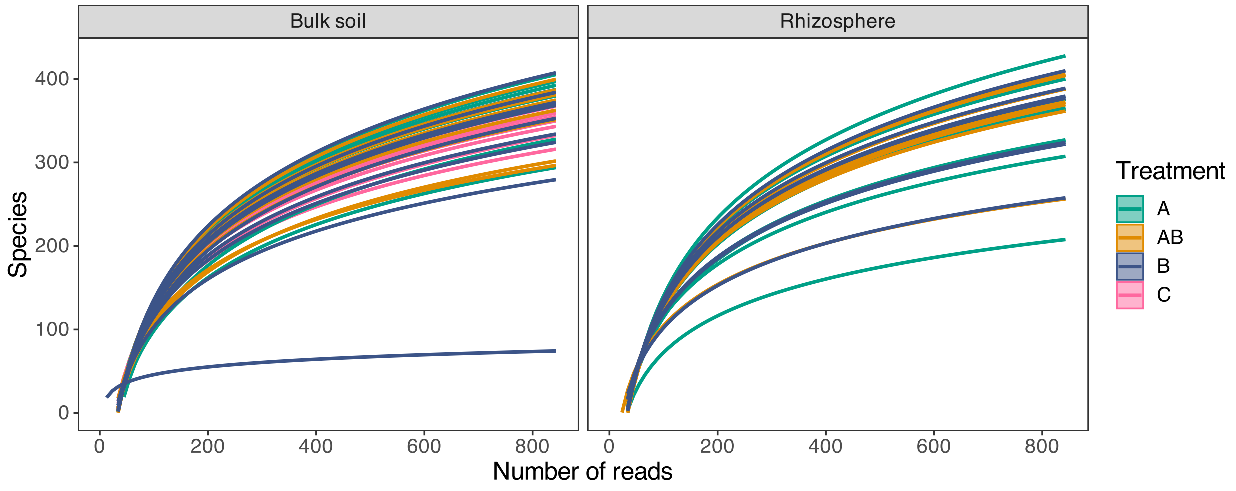

**Figure S1:** Rarefaction curves for 16S rRNA in soil samples. The curves illustrate rarefaction per soil type categorized by treatment (allium ’A’ samples are colored in green, banana ’B’ samples in blue, the co-cultivation allium + banana ’AB’ in yellow and the control ’C’ in pink).

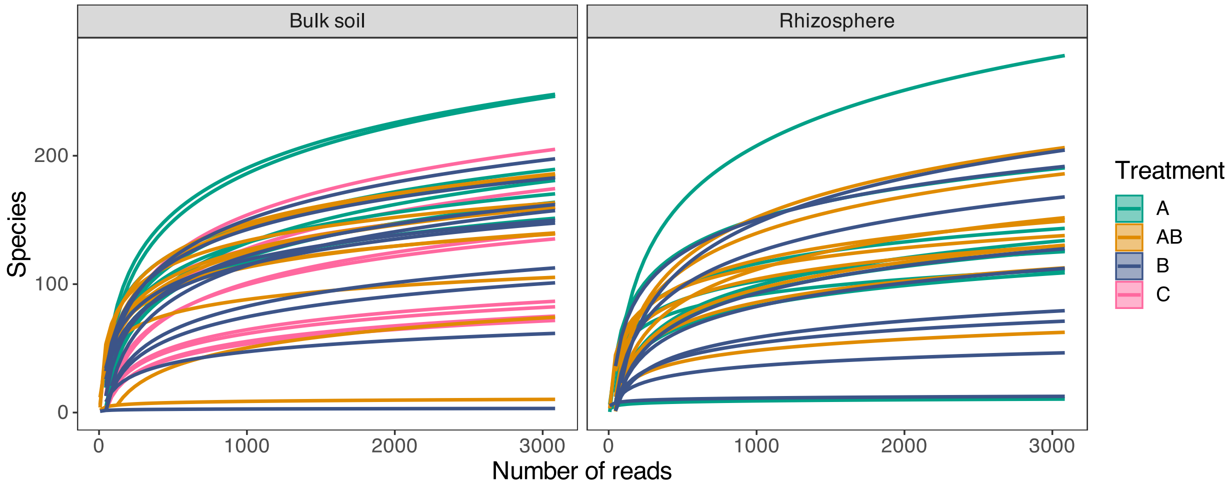

**Figure S2:** Rarefaction curves for ITS1 in soil samples. The curves illustrate rarefaction per soil type categorized by treatment (allium 'A' samples are colored in green, banana 'B' samples in blue, the co-cultivation allium + banana 'AB' in yellow and the control 'C' in pink).

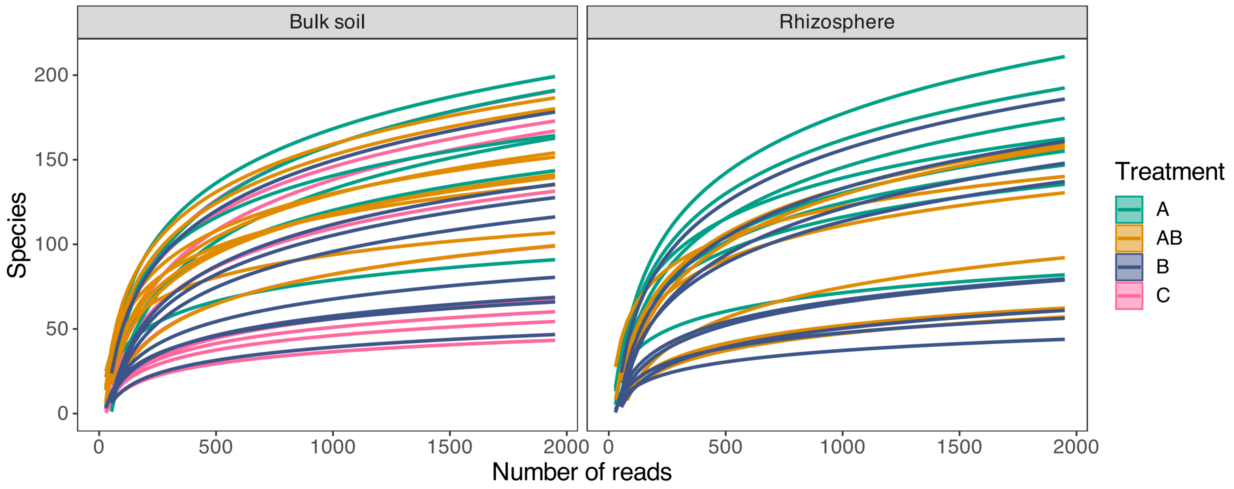

**Figure S3:** Rarefaction curves for ITS2 in soil samples. The curves illustrate rarefaction per soil type categorized by treatment (allium 'A' samples are colored in green, banana 'B' samples in blue, the co-cultivation allium + banana 'AB' in yellow and the control 'C' in pink).

**Table S1**: Abundance of bacterial (16S rRNA) phyla in bulk soil (Sb) and rhizosphere(Sr) categorized by treatment (allium 'A', banana 'B', co-cultivation 'AB' and control 'C'). Total (Tot), Rarefied (Abundance), and Rarefied Relative (Rel-Ab) Abundances as sequence counts are reported.

| Soil type | Phylum | A | | | AB | | | B | | | C | | |
|---|---|---|---|---|---|---|---|---|---|---|---|---|---|
| | | Total | Ab | Rel-Ab | Total | Ab | Rel-Ab | Total | Ab | Rel-Ab | Total | Ab | Rel-Ab |
| Sb | Acidobacteriota | 4761 | 529.00 ± 180.26 | 9.02 ± 3.08 | 5827 | 728.38 ± 252.68 | 12.42 ± 4.31 | 6715 | 671.50 ± 137.49 | 11.45 ± 2.35 | 5624 | 562.40 ± 77.62 | 9.59 ± 1.32 |
| | Actinobacteriota | 9812 | 1090.22 ± 124.03 | 18.6 ± 2.12 | 7493 | 936.62 ± 154.28 | 15.97 ± 2.63 | 10851 | 1085.10 ± 198.78 | 18.51 ± 3.39 | 12434 | 1243.40 ± 211.95 | 21.21 ± 3.61 |
| | Armatimonadota | 48 | 5.33 ± 3.32 | 0.09 ± 0.06 | 33 | 4.12 ± 3.27 | 0.07 ± 0.06 | 31 | 3.10 ± 2.02 | 0.05 ± 0.04 | 37 | 3.70 ± 2.45 | 0.06 ± 0.04 |
| | Bacteroidota | 2979 | 331.00 ± 51.83 | 5.65 ± 0.89 | 3782 | 472.75 ± 178.27 | 8.06 ± 3.04 | 3437 | 343.70 ± 84.47 | 5.86 ± 1.44 | 3845 | 384.50 ± 139.25 | 6.56 ± 2.37 |
| | Bdellovibrionota | 102 | 11.33 ± 16.16 | 0.20 ± 0.27 | 33 | 4.12 ± 3.18 | 0.07 ± 0.06 | 66 | 6.60 ± 6.87 | 0.11 ± 0.12 | 42 | 4.20 ± 3.08 | 0.07 ± 0.05 |
| | Campylobacterota | 35 | 3.89 ± 2.32 | 0.06 ± 0.04 | 51 | 6.38 ± 5.93 | 0.11 ± 0.10 | 67 | 6.70 ± 5.62 | 0.12 ± 0.10 | 91 | 9.10 ± 8.33 | 0.16 ± 0.14 |
| | Chloroflexi | 4023 | 447.00 ± 100.10 | 7.63 ± 1.71 | 4501 | 562.62 ± 175.5 | 9.60 ± 2.99 | 5206 | 520.6 ± 109.31 | 8.88 ± 1.86 | 4762 | 476.20 ± 130.11 | 8.12 ± 2.22 |
| | Cyanobacteria | 5023 | 558.11 ± 656.92 | 9.52 ± 11.21 | 1946 | 243.25 ± 666.74 | 4.15 ± 11.37 | 2940 | 294.00 ± 501.09 | 5.02 ± 8.55 | 2099 | 209.90 ± 374.51 | 3.58 ± 6.39 |
| | Dadabacteria | 65 | 7.22 ± 3.87 | 0.12 ± 0.07 | 131 | 16.38 ± 7.98 | 0.28 ± 0.14 | 132 | 13.20 ± 4.92 | 0.22 ± 0.08 | 79 | 7.90 ± 3.87 | 0.14 ± 0.07 |
| | Dependentiae | 119 | 13.22 ± 5.76 | 0.23 ± 0.10 | 73 | 9.12 ± 4.67 | 0.16 ± 0.08 | 81 | 8.10 ± 5.07 | 0.14 ± 0.09 | 106 | 10.60 ± 3.98 | 0.18 ± 0.07 |
| | Desulfobacterota | 16 | 1.78 ± 5.33 | 0.03 ± 0.09 | 4 | 0.50 ± 1.41 | 0.01 ± 0.02 | 21 | 2.10 ± 5.67 | 0.04 ± 0.10 | 2 | 0.20 ± 0.63 | 0.00 ± 0.01 |
| | Elusimicrobiota | 16 | 1.78 ± 1.48 | 0.03 ± 0.02 | 25 | 3.12 ± 3.04 | 0.05 ± 0.05 | 31 | 3.10 ± 3.07 | 0.05 ± 0.05 | 19 | 1.90 ± 3.11 | 0.03 ± 0.05 |
| | Entotheonellaeota | 40 | 4.44 ± 2.65 | 0.08 ± 0.04 | 47 | 5.88 ± 2.42 | 0.10 ± 0.04 | 38 | 3.80 ± 2.74 | 0.06 ± 0.05 | 51 | 5.10 ± 2.77 | 0.09 ± 0.05 |
| | Fibrobacterota | 25 | 2.78 ± 1.64 | 0.05 ± 0.03 | 14 | 1.75 ± 1.98 | 0.03 ± 0.03 | 36 | 3.60 ± 4.70 | 0.06 ± 0.08 | 48 | 4.80 ± 3.29 | 0.08 ± 0.06 |
| | Firmicutes | 2675 | 297.22 ± 75.42 | 5.07 ± 1.29 | 2586 | 323.25 ± 95.49 | 5.51 ± 1.63 | 4763 | 476.30 ± 222.42 | 8.12 ± 3.79 | 3410 | 341.00 ± 179.21 | 5.82 ± 3.06 |
| | Fusobacteriota | 302 | 33.56 ± 24.34 | 0.57 ± 0.42 | 387 | 48.38 ± 21.27 | 0.82 ± 0.36 | 470 | 47.00 ± 17.64 | 0.80 ± 0.30 | 547 | 54.70 ± 34.90 | 0.93 ± 0.60 |
| | Gemmatimonadota | 1164 | 129.33 ± 35.06 | 2.21 ± 0.60 | 1134 | 141.75 ± 33.8 | 2.42 ± 0.58 | 1252 | 125.20 ± 26.62 | 2.14 ± 0.45 | 1273 | 127.30 ± 28.54 | 2.17 ± 0.49 |
| | Hydrogenedentes | 14 | 1.56 ± 1.88 | 0.03 ± 0.03 | 40 | 5.00 ± 2.88 | 0.09 ± 0.05 | 40 | 4.00 ± 3.20 | 0.07 ± 0.05 | 10 | 1.00 ± 1.05 | 0.02 ± 0.02 |
| | Methylomirabilota | 58 | 6.44 ± 4.77 | 0.11 ± 0.08 | 88 | 11.00 ± 6.05 | 0.19 ± 0.10 | 72 | 7.20 ± 5.39 | 0.12 ± 0.09 | 75 | 7.50 ± 3.60 | 0.13 ± 0.06 |
| | Myxococcota | 138 | 15.33 ± 7.87 | 0.26 ± 0.13 | 130 | 16.25 ± 10.89 | 0.28 ± 0.18 | 135 | 13.50 ± 11.54 | 0.23 ± 0.20 | 109 | 10.90 ± 7.64 | 0.19 ± 0.13 |
| | Nitrospirota | 56 | 6.22 ± 3.31 | 0.11 ± 0.06 | 51 | 6.38 ± 3.34 | 0.11 ± 0.06 | 34 | 3.40 ± 3.60 | 0.06 ± 0.06 | 78 | 7.80 ± 5.96 | 0.13 ± 0.10 |
| | Patescibacteria | 275 | 30.56 ± 11.44 | 0.52 ± 0.20 | 156 | 19.50 ± 9.62 | 0.33 ± 0.16 | 229 | 22.90 ± 14.62 | 0.39 ± 0.25 | 247 | 24.70 ± 12.14 | 0.42 ± 0.21 |
| | Planctomycetota | 1076 | 119.56 ± 52.06 | 2.04 ± 0.89 | 1083 | 135.38 ± 58.34 | 2.31 ± 1.00 | 1033 | 103.30 ± 43.60 | 1.76 ± 0.74 | 1160 | 116.00 ± 42.06 | 1.98 ± 0.72 |
| | Proteobacteria | 19821 | 2202.33 ± 374.77 | 37.56 ± 6.39 | 17174 | 2146.75 ± 218.28 | 36.62 ± 3.73 | 20762 | 2076.20 ± 255.88 | 35.41 ± 4.36 | 22288 | 2228.80 ± 127.73 | 38.01 ± 2.18 |
| | Spirochaetota | 6 | 0.67 ± 2.00 | 0.01 ± 0.03 | 3 | 0.38 ± 0.74 | 0.01 ± 0.01 | 5 | 0.50 ± 1.58 | 0.01 ± 0.03 | 0 | 0.00 ± 0.00 | 0.00 ± 0.00 |
| | Sumerlaeota | 0 | 0.00 ± 0.00 | 0.00 ± 0.00 | 0 | 0.00 ± 0.00 | 0.00 ± 0.00 | 0 | 0.00 ± 0.00 | 0.00 ± 0.00 | 1 | 0.10 ± 0.32 | 0.00 ± 0.01 |
| | Verrucomicrobiota | 118 | 13.11 ± 3.82 | 0.22 ± 0.07 | 112 | 14.00 ± 6.07 | 0.24 ± 0.10 | 183 | 18.30 ± 6.41 | 0.31 ± 0.11 | 190 | 19.00 ± 7.83 | 0.33 ± 0.13 |
| | WS2 | 0 | 0.00 ± 0.00 | 0.00 ± 0.00 | 0 | 0.00 ± 0.00 | 0.00 ± 0.00 | 0 | 0.00 ± 0.00 | 0.00 ± 0.00 | 3 | 0.30 ± 0.67 | 0.00 ± 0.01 |
| Sr | Abditibacteriota | 0 | 0.00 ± 0.00 | 0.00 ± 0.00 | 0 | 0.00 ± 0.00 | 0.00 ± 0.00 | 1 | 0.14 ± 0.38 | 0.02 ± 0.04 | - | - | - |
| | Acidobacteriota | 1027 | 114.11 ± 31.31 | 12.54 ± 3.44 | 538 | 76.86 ± 31.76 | 8.44 ± 3.49 | 397 | 56.71 ± 15.14 | 6.23 ± 1.66 | - | - | - |
| | Actinobacteriota | 1617 | 179.67 ± 54.67 | 19.74 ± 6.01 | 912 | 130.29 ± 22.35 | 14.32 ± 2.45 | 967 | 138.14 ± 18.72 | 15.18 ± 2.05 | - | - | - |
| | Armatimonadota | 3 | 0.33 ± 0.50 | 0.04 ± 0.05 | 9 | 1.29 ± 2.36 | 0.14 ± 0.26 | 17 | 2.43 ± 1.81 | 0.27 ± 0.20 | - | - | - |
| | Bacteroidota | 426 | 47.33 ± 14.83 | 5.20 ± 1.63 | 238 | 34.00 ± 10.97 | 3.74 ± 1.21 | 322 | 46.00 ± 15.50 | 5.06 ± 1.70 | - | - | - |
| | Bdellovibrionota | 13 | 1.44 ± 1.94 | 0.16 ± 0.21 | 29 | 4.14 ± 3.63 | 0.46 ± 0.40 | 25 | 3.57 ± 2.57 | 0.39 ± 0.28 | - | - | - |
| | Campylobacterota | 6 | 0.67 ± 0.71 | 0.07 ± 0.08 | 2 | 0.29 ± 0.49 | 0.03 ± 0.05 | 6 | 0.86 ± 1.21 | 0.09 ± 0.13 | - | - | - |
| | Chloroflexi | 776 | 86.22 ± 15.43 | 9.47 ± 1.70 | 456 | 65.14 ± 18.37 | 7.16 ± 2.02 | 347 | 49.57 ± 9.71 | 5.44 ± 1.07 | - | - | - |
| | Cyanobacteria | 1 | 0.11 ± 0.33 | 0.01 ± 0.04 | 18 | 2.57 ± 2.23 | 0.28 ± 0.24 | 17 | 2.43 ± 2.37 | 0.27 ± 0.26 | - | - | - |
| | Dadabacteria | 20 | 2.22 ± 1.56 | 0.24 ± 0.17 | 7 | 1.00 ± 1.41 | 0.11 ± 0.16 | 5 | 0.71 ± 0.95 | 0.08 ± 0.10 | - | - | - |
| | Dependentiae | 21 | 2.33 ± 2.29 | 0.26 ± 0.25 | 14 | 2.00 ± 3.27 | 0.22 ± 0.36 | 6 | 0.86 ± 1.07 | 0.09 ± 0.12 | - | - | - |
| | Desulfobacterota | 2 | 0.22 ± 0.67 | 0.02 ± 0.07 | 7 | 1.00 ± 2.65 | 0.11 ± 0.29 | 5 | 0.71 ± 1.89 | 0.08 ± 0.21 | - | - | - |
| | Elusimicrobiota | 2 | 0.22 ± 0.67 | 0.02 ± 0.07 | 0 | 0.00 ± 0.00 | 0.00 ± 0.00 | 0 | 0.00 ± 0.00 | 0.00 ± 0.00 | - | - | - |
| | Entotheonellaeota | 3 | 0.33 ± 1.00 | 0.04 ± 0.11 | 2 | 0.29 ± 0.76 | 0.03 ± 0.08 | 1 | 0.14 ± 0.38 | 0.02 ± 0.04 | - | - | - |
| | Fibrobacterota | 8 | 0.89 ± 1.05 | 0.10 ± 0.12 | 7 | 1.00 ± 1.41 | 0.11 ± 0.16 | 3 | 0.43 ± 0.53 | 0.05 ± 0.06 | - | - | - |

| | | | | | | | | | | | | | |
|---|---|---|---|---|---|---|---|---|---|---|---|---|---|
| | Firmicutes | 912 | 101.33 ± 47.41 | 11.13 ± 5.21 | 998 | 142.57 ± 89.51 | 15.67 ± 9.84 | 1035 | 147.86 ± 48.69 | 16.25 ± 5.35 | - | - | - |
| | Fusobacteriota | 42 | 4.67 ± 2.60 | 0.51 ± 0.29 | 28 | 4.00 ± 3.79 | 0.44 ± 0.42 | 26 | 3.71 ± 2.87 | 0.41 ± 0.32 | - | - | - |
| | Gemmatimonadota | 183 | 20.33 ± 7.91 | 2.24 ± 0.87 | 91 | 13.00 ± 7.07 | 1.43 ± 0.78 | 93 | 13.29 ± 4.15 | 1.46 ± 0.46 | - | - | - |
| | Hydrogenedentes | 3 | 0.33 ± 0.50 | 0.04 ± 0.06 | 1 | 0.14 ± 0.38 | 0.02 ± 0.04 | 0 | 0.00 ± 0.00 | 0.00 ± 0.00 | - | - | - |
| | Methylomirabilota | 9 | 1.00 ± 0.87 | 0.11 ± 0.10 | 8 | 1.14 ± 1.35 | 0.13 ± 0.15 | 3 | 0.43 ± 0.53 | 0.05 ± 0.06 | - | - | - |
| | Myxococcota | 27 | 3.00 ± 4.61 | 0.33 ± 0.51 | 82 | 11.71 ± 14.19 | 1.29 ± 1.56 | 76 | 10.86 ± 5.98 | 1.19 ± 0.66 | - | - | - |
| | Nitrospirota | 6 | 0.67 ± 0.87 | 0.07 ± 0.10 | 1 | 0.14 ± 0.38 | 0.02 ± 0.04 | 3 | 0.43 ± 0.53 | 0.05 ± 0.06 | - | - | - |
| | Patescibacteria | 30 | 3.33 ± 1.41 | 0.37 ± 0.16 | 75 | 10.71 ± 7.3 | 1.18 ± 0.80 | 94 | 13.43 ± 7.44 | 1.48 ± 0.82 | - | - | - |
| | Planctomycetota | 150 | 16.67 ± 5.74 | 1.83 ± 0.63 | 83 | 11.86 ± 6.69 | 1.30 ± 0.74 | 68 | 9.71 ± 3.30 | 1.07 ± 0.36 | - | - | - |
| | Proteobacteria | 2881 | 320.11 ± 43.78 | 35.18 ± 4.81 | 2730 | 390.00 ± 88.72 | 42.86 ± 9.75 | 2814 | 402.00 ± 80.11 | 44.18 ± 8.80 | - | - | - |
| | Verrucomicrobiota | 20 | 2.22 ± 0.97 | 0.24 ± 0.11 | 34 | 4.86 ± 3.29 | 0.53 ± 0.36 | 38 | 5.43 ± 2.15 | 0.60 ± 0.24 | - | - | - |
| | WS2 | 2 | 0.22 ± 0.67 | 0.02 ± 0.07 | 0 | 0.00 ± 0.00 | 0.00 ± 0.00 | 1 | 0.14 ± 0.38 | 0.02 ± 0.04 | - | - | - |

**Table S2**: Abundance of fungal (ITS1 and ITS2) phyla in bulk soil (Sb) and rhizosphere (Sr) categorized by treatment (allium 'A', banana 'B', co-cultivation 'AB' and control 'C'). Total (Tot), Rarefied (Ab), and Rarefied Relative (Rel-Ab) Abundance sequence counts are reported.

| **ITS** | **Soil type** | **Phylum** | **A** | | | **AB** | | | **B** | | | **C** | | |
|---|---|---|---|---|---|---|---|---|---|---|---|---|---|---|
| | | | Total | Ab | Rel-Ab | Total | Ab | Rel-Ab | Total | Ab | Rel-Ab | Total | Ab | Rel-Ab |
| **1** | Sb | Aphelidiomycota | 0 | 0.00 ± 0.00 | 0.00 ± 0.00 | 2 | 0.20 ± 0.42 | 0.16 ± 0.33 | 1 | 0.1 ± 0.32 | 0.08 ± 0.25 | 1 | 0.10 ± 0.32 | 0.08 ± 0.25 |
| | | Ascomycota | 869 | 108.62 ± 6.93 | 86.21 ± 5.50 | 976 | 97.60 ± 16.24 | 77.46 ± 12.89 | 1054 | 105.4 ± 9.91 | 83.65 ± 7.87 | 1038 | 103.80 ± 36.75 | 82.38 ± 29.17 |
| | | Basidiomycota | 73 | 9.12 ± 4.42 | 7.24 ± 3.51 | 137 | 13.70 ± 7.69 | 10.87 ± 6.10 | 97 | 9.7 ± 6.72 | 7.7 ± 5.33 | 171 | 17.10 ± 37.62 | 13.57 ± 29.86 |
| | | Chytridiomycota | 8 | 1.00 ± 0.76 | 0.79 ± 0.60 | 8 | 0.80 ± 0.92 | 0.64 ± 0.73 | 7 | 0.7 ± 1.06 | 0.56 ± 0.84 | 8 | 0.80 ± 1.32 | 0.63 ± 1.04 |
| | | Mortierellomycota | 23 | 2.88 ± 3.68 | 2.28 ± 2.92 | 84 | 8.40 ± 10.25 | 6.67 ± 8.14 | 78 | 7.8 ± 8.19 | 6.19 ± 6.5 | 22 | 2.20 ± 1.75 | 1.74 ± 1.39 |
| | | Mucoromycota | 22 | 2.75 ± 2.66 | 2.18 ± 2.11 | 33 | 3.30 ± 3.33 | 2.62 ± 2.65 | 18 | 1.8 ± 2.15 | 1.43 ± 1.71 | 7 | 0.70 ± 0.48 | 0.55 ± 0.38 |
| | | Rozellomycota | 13 | 1.62 ± 2.72 | 1.29 ± 2.16 | 20 | 2.00 ± 2.05 | 1.59 ± 1.63 | 5 | 0.5 ± 0.85 | 0.4 ± 0.68 | 13 | 1.30 ± 1.95 | 1.03 ± 1.54 |
| | Sr | Aphelidiomycota | 34 | 3.78 ± 3.90 | 0.12 ± 0.13 | 6 | 0.67 ± 0.71 | 0.02 ± 0.02 | 5 | 0.56 ± 0.73 | 0.02 ± 0.02 | - | - | - |
| | | Ascomycota | 20880 | 2320.00 ± 271.93 | 75.23 ± 8.82 | 24456 | 2717.33 ± 422.36 | 88.11 ± 13.70 | 26080 | 2897.78 ± 90.74 | 93.96 ± 2.94 | - | - | - |
| | | Basidiobolomycota | 70 | 7.78 ± 17.16 | 0.25 ± 0.56 | 8 | 0.89 ± 2.67 | 0.03 ± 0.09 | 0 | 0.00 ± 0.00 | 0.00 ± 0.00 | - | - | - |
| | | Basidiomycota | 4398 | 488.67 ± 343.36 | 15.85 ± 11.13 | 2602 | 289.11 ± 309.49 | 9.37 ± 10.03 | 1447 | 160.78 ± 71.24 | 5.21 ± 2.31 | - | - | - |
| | | Chytridiomycota | 172 | 19.11 ± 28.94 | 0.62 ± 0.94 | 175 | 19.44 ± 40.69 | 0.63 ± 1.32 | 35 | 3.89 ± 3.10 | 0.12 ± 0.10 | - | - | - |
| | | Glomeromycota | 28 | 3.11 ± 8.27 | 0.10 ± 0.27 | 0 | 0.00 ± 0.00 | 0.00 ± 0.00 | 4 | 0.44 ± 0.73 | 0.01 ± 0.02 | - | - | - |
| | | Kickxellomycota | 18 | 2.00 ± 4.24 | 0.06 ± 0.14 | 14 | 1.56 ± 4.67 | 0.05 ± 0.15 | 3 | 0.33 ± 1.00 | 0.01 ± 0.03 | - | - | - |
| | | Mortierellomycota | 1537 | 170.78 ± 177.67 | 5.54 ± 5.76 | 104 | 11.56 ± 19.93 | 0.37 ± 0.65 | 123 | 13.67 ± 26.05 | 0.44 ± 0.85 | - | - | - |
| | | Mucoromycota | 368 | 40.89 ± 43.07 | 1.33 ± 1.40 | 237 | 26.33 ± 65.21 | 0.85 ± 2.11 | 11 | 1.22 ± 2.05 | 0.04 ± 0.07 | - | - | - |
| | | Olpidiomycota | 0 | 0.00 ± 0.00 | 0.00 ± 0.00 | 0 | 0.00 ± 0.00 | 0.00 ± 0.00 | 1 | 0.11 ± 0.33 | 0 .00± 0.01 | - | - | - |
| | | Rozellomycota | 251 | 27.89 ± 27.06 | 0.90 ± 0.88 | 154 | 17.11 ± 29.76 | 0.55 ± 0.97 | 47 | 5.22 ± 3.96 | 0.17 ± 0.13 | - | - | - |
| **2** | Sb | Aphelidiomycota | 90 | 15.00 ± 8.00 | 0.76 ± 0.41 | 215 | 23.89 ± 12.22 | 1.22 ± 0.63 | 83 | 10.38 ± 12.98 | 0.53 ± 0.67 | 38 | 4.22 ± 1.48 | 0.21 ± 0.08 |
| | | Ascomycota | 9872 | 1645.33 ± 151.74 | 84.16 ± 7.76 | 13935 | 1548.33 ± 112.46 | 79.20 ± 5.75 | 13758 | 1719.75 ± 176.22 | 87.97 ± 9.02 | 15040 | 1671.11 ± 201.99 | 85.48 ± 10.33 |
| | | Basidiobolomycota | 4 | 0.67 ± 1.21 | 0.03 ± 0.06 | 0 | 0.00 ± 0.00 | 0.00 ± 0.00 | 0 | 0.00 ± 0.00 | 0.00 ± 0.00 | 2 | 0.22 ± 0.67 | 0.01 ± 0.03 |
| | | Basidiomycota | 1168 | 194.67 ± 114.99 | 9.96 ± 5.88 | 1945 | 216.11 ± 91.80 | 11.06 ± 4.70 | 899 | 112.38 ± 66.21 | 5.75 ± 3.39 | 1300 | 144.44 ± 76.21 | 7.39 ± 3.90 |
| | | Chytridiomycota | 107 | 17.83 ± 7.94 | 0.91 ± 0.41 | 212 | 23.56 ± 30.55 | 1.20 ± 1.56 | 53 | 6.62 ± 3.89 | 0.34 ± 0.20 | 731 | 81.22 ± 212.13 | 4.15 ± 10.85 |
| | | Mortierellomycota | 139 | 23.17 ± 27.69 | 1.19 ± 1.42 | 619 | 68.78 ± 100.93 | 3.52 ± 5.16 | 642 | 80.25 ± 147.93 | 4.11 ± 7.57 | 216 | 24.00 ± 19.43 | 1.23 ± 0.99 |
| | | Mucoromycota | 139 | 23.17 ± 18.63 | 1.19 ± 0.95 | 248 | 27.56 ± 16.82 | 1.41 ± 0.86 | 122 | 15.25 ± 18.20 | 0.78 ± 0.93 | 82 | 9.11 ± 7.24 | 0.46 ± 0.37 |
| | | Rozellomycota | 211 | 35.17 ± 35.74 | 1.80 ± 1.83 | 421 | 46.78 ± 30.59 | 2.39 ± 1.57 | 83 | 10.38 ± 16.96 | 0.53 ± 0.87 | 186 | 20.67 ± 12.47 | 1.06 ± 0.64 |
| | Sr | Aphelidiomycota | 230 | 28.75 ± 26.85 | 0.62 ± 0.58 | 73 | 10.43 ± 12.55 | 0.22 ± 0.27 | 94 | 10.44 ± 11.67 | 0.22 ± 0.25 | - | - | - |
| | | Ascomycota | 30850 | 3856.25 ± 230.86 | 82.57 ± 4.94 | 30205 | 4315.00 ± 226.69 | 92.40 ± 4.86 | 38267 | 4251.89 ± 382.83 | 91.05 ± 8.20 | - | - | - |
| | | Basidiobolomycota | 0 | 0.00 ± 0.00 | 0.00 ± 0.00 | 0 | 0.00 ± 0.00 | 0.00 ± 0.00 | 8 | 0.89 ± 2.03 | 0.02 ± 0.04 | - | - | - |
| | | Basidiomycota | 4224 | 528.00 ± 208.66 | 11.30 ± 4.47 | 1776 | 253.71 ± 133.92 | 5.43 ± 2.87 | 2588 | 287.56 ± 190.07 | 6.16 ± 4.07 | - | - | - |
| | | Chytridiomycota | 120 | 15.00 ± 20.40 | 0.32 ± 0.44 | 60 | 8.57 ± 12.42 | 0.18 ± 0.27 | 95 | 10.56 ± 9.59 | 0.22 ± 0.20 | - | - | - |
| | | Mortierellomycota | 1320 | 165.00 ± 128.33 | 3.54 ± 2.75 | 197 | 28.14 ± 69.68 | 0.60 ± 1.49 | 705 | 78.33 ± 149.09 | 1.68 ± 3.19 | - | - | - |
| | | Mucoromycota | 298 | 37.25 ± 23.97 | 0.80 ± 0.51 | 141 | 20.14 ± 24.64 | 0.43 ± 0.53 | 135 | 15.00 ± 15.95 | 0.32 ± 0.34 | - | - | - |
| | | Rozellomycota | 318 | 39.75 ± 22.19 | 0.85 ± 0.48 | 238 | 34.00 ± 53.88 | 0.73 ± 1.16 | 138 | 15.33 ± 23.09 | 0.33 ± 0.49 | - | - | - |

**Table S3:** Summary of differentially abundant and linear discriminant analysis (LDA) for bacterial genera in (Sb) and rhizosphere (Sr). ’Tx’ indicates the treatment in which the ASV exhibited significant abundance (allium ’A’, banana ’B’, co-cultivation ’AB’ and control ’C’). ’FDR’ is false discovery rate.

| **Soil type** | **Taxonomy** | | | | | **LDA upper** | **LDA mean** | **LDA lower** | **Tx** | **p - Value** | **FDR** |
|---|---|---|---|---|---|---|---|---|---|---|---|
| | **Phylum** | **Class** | **Order** | **Family** | **Genus** | | | | | | |
| **Sb** | Cyanobacteria | Cyanobacteriia | Geitlerinematales | Geitlerinemataceae | Geitlerinema | 4.121 | 4.055 | 3.977 | A | 0.000 | 0.000 |
| | Cyanobacteria | Cyanobacteriia | Phormidesmiales | Nodosilineaceae | Nodosilinea PCC-7104 | 2.433 | 2.383 | 2.327 | A | 0.000 | 0.022 |
| | Planctomycetota | Planctomycetes | Planctomycetales | Schlesneriaceae | Planctopirus | 2.402 | 2.357 | 2.306 | AB | 0.000 | 0.017 |
| | Proteobacteria | Gammaproteobacteria | Burkholderiales | Comamonadaceae | Rhizobacter | 2.481 | 2.417 | 2.342 | B | 0.000 | 0.027 |
| | Actinobacteriota | Actinobacteria | Micromonosporales | Micromonosporaceae | Rhizocola | 2.409 | 2.365 | 2.316 | A | 0.000 | 0.027 |
| **Sr** | Proteobacteria | Gammaproteobacteria | Burkholderiales | Comamonadaceae | Aquabacterium | 3.278 | 3.226 | 3.166 | B | 0.000 | 0.113 |
| | Proteobacteria | Alphaproteobacteria | Caulobacterales | Caulobacteraceae | Asticcacaulis | 3.181 | 3.131 | 3.074 | B | 0.000 | 0.127 |
| | Patescibacteria | ABY1 | Candidatus Kuenenbacteria | Candidatus Kuenenbacteria | Candidatus Kuenenbacteria | 3.097 | 3.044 | 2.985 | A | 0.003 | 0.147 |
| | Myxococcota | Myxococcia | Myxococcales | Myxococcaceae | Corallococcus | 3.171 | 3.123 | 3.069 | B | 0.005 | 0.175 |
| | Actinobacteriota | Acidimicrobiia | Microtrichales | Ilumatobacteraceae | Ilumatobacter | 3.276 | 3.228 | 3.174 | A | 0.006 | 0.178 |
| | Proteobacteria | Gammaproteobacteria | Burkholderiales | Nitrosomonadaceae | IS-44 | 2.999 | 2.940 | 2.872 | A | 0.008 | 0.204 |
| | Acidobacteriota | Blastocatellia | Pyrinomonadales | Pyrinomonadaceae | RB41 | 3.819 | 3.777 | 3.731 | A | 0.006 | 0.179 |
| | Proteobacteria | Gammaproteobacteria | Burkholderiales | TRA3-20 | TRA3-20 | 3.420 | 3.368 | 3.310 | A | 0.001 | 0.147 |

**Table S4:** Abundance of bacterial (16S rRNA) differentially abundant genera in bulk soil (Sb) and rhizosphere (Sr) categorized by treatment (allium 'A', banana 'B', co- cultivation 'AB' and control 'C'). Rarefied (Ab), and Rarefied Relative (Rel-Ab) Abundances are reported.

| Soil type | Genus | A | | AB | | B | | C | |
|---|---|---|---|---|---|---|---|---|---|
| | | Ab | Rel-Ab | Ab | Rel-Ab | Ab | Rel-Ab | Ab | Rel-Ab |
| **Sb** | Geitlerinema | 258.44 ± 447.54 | 4.41 ± 7.63 | 1.00 ± 2.83 | 0.02 ± 0.05 | 0.00 ± 0.00 | 0.00 ± 0.00 | 0.00 ± 0.00 | 0.00 ± 0.00 |
| | Nodosilinea PCC-7104 | 5.22 ± 4.32 | 0.09 ± 0.07 | 0.00 ± 0.00 | 0.00 ± 0.00 | 1.10 ± 2.60 | 0.02 ± 0.04 | 0.00 ± 0.00 | 0.00 ± 0.00 |
| | Planctopirus | 0.11 ± 0.33 | 0.00 ± 0.01 | 4.75 ± 2.82 | 0.08 ± 0.05 | 1.20 ± 1.62 | 0.02 ± 0.03 | 0.00 ± 0.00 | 0.00 ± 0.00 |
| | Rhizobacter | 0.00 ± 0.00 | 0.00 ± 0.00 | 0.00 ± 0.00 | 0.00 ± 0.00 | 6.50 ± 9.12 | 0.11 ± 0.16 | 0.00 ± 0.00 | 0.00 ± 0.00 |
| | Rhizocola | 5.22 ± 3.23 | 0.09 ± 0.06 | 0.38 ± 1.06 | 0.01 ± 0.02 | 0.30 ± 0.95 | 0.01 ± 0.02 | 1.50 ± 1.78 | 0.03 ± 0.03 |
| **Sr** | Aquabacterium | 0.00 ± 0.00 | 0.00 ± 0.00 | 0.14 ± 0.38 | 0.02 ± 0.04 | 3.71 ± 2.29 | 0.41 ± 0.25 | - | - |
| | Asticcacaulis | 0.00 ± 0.00 | 0.00 ± 0.00 | 0.57 ± 0.98 | 0.06 ± 0.11 | 2.29 ± 1.25 | 0.25 ± 0.14 | - | - |
| | Candidatus Kuenenbacteria | 1.56 ± 1.01 | 0.17 ± 0.11 | 0.29 ± 0.49 | 0.03 ± 0.05 | 0.00 ± 0.00 | 0.00 ± 0.00 | - | - |
| | Corallococcus | 0.00 ± 0.00 | 0.00 ± 0.00 | 0.29 ± 0.49 | 0.03 ± 0.05 | 2.29 ± 2.14 | 0.25 ± 0.23 | - | - |
| | Ilumatobacter | 5.22 ± 2.11 | 0.57 ± 0.23 | 2.29 ± 1.80 | 0.25 ± 0.20 | 1.71 ± 1.11 | 0.19 ± 0.12 | - | - |
| | IS-44 | 1.22 ± 1.09 | 0.13 ± 0.12 | 0.14 ± 0.38 | 0.02 ± 0.04 | 0.00 ± 0.00 | 0.00 ± 0.00 | - | - |
| | RB41 | 19.78 ± 10.10 | 2.17 ± 1.11 | 9.29 ± 11.00 | 1.02 ± 1.21 | 4.14 ± 2.27 | 0.46 ± 0.25 | - | - |
| | TRA3-20 | 7.56 ± 2.51 | 0.83 ± 0.28 | 2.14 ± 2.19 | 0.24 ± 0.24 | 2.14 ± 2.27 | 0.24 ± 0.25 | - | - |

**Table S5:** Summary of differentially abundant and linear discriminant analysis (LDA) for fungal genera in (Sb) and rhizosphere (Sr). ’Tx’ indicates the treatment in which the ASV exhibited significant abundance (allium ’A’, banana ’B’, co-cultivation ’AB’ and control ’C’). ’FDR’ stands for false discovery rate.

| ITS | Soil type | Taxonomy | | | | | LDA upper | LDA mean | LDA lower | Tx | p - Value | FDR |
|---|---|---|---|---|---|---|---|---|---|---|---|---|
| | | **Phylum** | **Class** | **Order** | **Family** | **Genus** | | | | | | |
| **1** | **Sb** | Ascomycota | Sordariomycetes | Hypocreales | Incertae sedis | Acremonium | 4.848 | 4.794 | 4.733 | A | 0.035 | 0.596 |
| | | Ascomycota | Eurotiomycetes | Eurotiales | Aspergillaceae | Aspergillus | 3.516 | 3.461 | 3.397 | A | 0.015 | 0.596 |
| | **Sr** | Ascomycota | Sordariomycetes | Sordariales | Lasiosphaeriaceae | Apodus | 2.744 | 2.693 | 2.636 | B | 0.012 | 0.360 |
| | | Ascomycota | Pezizomycetes | Pezizales | Ascobolaceae | Ascobolus | 3.856 | 3.801 | 3.738 | B | 0.019 | 0.386 |
| | | Ascomycota | Dothideomycetes | Capnodiales | Capnodiaceae | Leptoxyphium | 2.705 | 2.617 | 2.507 | A | 0.001 | 0.313 |
| | | Mortierellomycota | Mortierellomycetes | Mortierellales | Mortierellaceae | Mortierella | 4.232 | 4.170 | 4.097 | A | 0.020 | 0.386 |
| | | Basidiomycota | Tremellomycetes | Cystofilobasidiales | Mrakiaceae | Tausonia | 3.370 | 3.311 | 3.243 | B | 0.004 | 0.313 |
| | | Ascomycota | Sordariomycetes | Hypocreales | Hypocreaceae | Trichoderma | 3.370 | 3.314 | 3.251 | A | 0.047 | 0.454 |
| **2** | **Sb** | Ascomycota | Sordariomycetes | Microascales | Microascaceae | Scedosporium | 3.882 | 3.837 | 3.786 | AB | 0.002 | 0.162 |
| | | Ascomycota | Sordariomycetes | Glomerellales | Plectosphaerellaceae | Wallrothiella | 3.108 | 3.059 | 3.004 | AB | 0.004 | 0.171 |
| | **Sr** | Basidiomycota | Tremellomycetes | Trichosporonales | Trichosporonaceae | Apiotrichum | 4.064 | 4.012 | 3.953 | A | 0.006 | 0.135 |
| | | Ascomycota | Sordariomycetes | Incertae sedis | Incertae sedis | Aquapteridospora | 2.865 | 2.801 | 2.727 | A | 0.006 | 0.135 |
| | | Ascomycota | Orbiliomycetes | Orbiliales | Orbiliaceae | Arthrobotrys | 3.440 | 3.382 | 3.315 | A | 0.045 | 0.232 |
| | | Ascomycota | Sordariomycetes | Chaetosphaeriales | Chaetosphaeriaceae | Chloridium | 2.977 | 2.909 | 2.828 | A | 0.010 | 0.135 |
| | | Basidiomycota | Agaricomycetes | Agaricales | Psathyrellaceae | Coprinopsis | 3.678 | 3.627 | 3.570 | A | 0.002 | 0.135 |
| | | Ascomycota | Sordariomycetes | Hypocreales | Nectriaceae | Cylindrocladiella | 2.437 | 2.367 | 2.284 | A | 0.011 | 0.135 |
| | | Ascomycota | Sordariomycetes | Hypocreales | Nectriaceae | Fusarium | 4.320 | 4.262 | 4.196 | A | 0.029 | 0.205 |
| | | Ascomycota | Leotiomycetes | Helotiales | Hyaloscyphaceae | Hyaloscypha | 2.558 | 2.486 | 2.399 | A | 0.010 | 0.135 |
| | | Ascomycota | Dothideomycetes | Botryosphaeriales | Botryosphaeriaceae | Lasiodiplodia | 2.976 | 2.903 | 2.815 | A | 0.008 | 0.135 |
| | | Ascomycota | Sordariomycetes | Microascales | Microascaceae | Microascus | 2.675 | 2.618 | 2.553 | A | 0.021 | 0.188 |
| | | Mortierellomycota | Mortierellomycetes | Mortierellales | Mortierellaceae | Mortierella | 3.929 | 3.884 | 3.833 | A | 0.008 | 0.135 |
| | | Ascomycota | Sordariomycetes | Sordariales | Chaetomiaceae | Mycothermus | 3.920 | 3.871 | 3.817 | A | 0.006 | 0.135 |
| | | Ascomycota | Sordariomycetes | Microascales | Microascaceae | Scedosporium | 4.137 | 4.076 | 4.005 | A | 0.002 | 0.135 |
| | | Basidiomycota | Tremellomycetes | Cystofilobasidiales | Mrakiaceae | Tausonia | 3.269 | 3.210 | 3.142 | B | 0.030 | 0.205 |
| | | Ascomycota | Sordariomycetes | Sordariales | Chaetomiaceae | Zopfiella | 3.381 | 3.322 | 3.253 | AB | 0.024 | 0.189 |

**Table S6:** Abundance of fungal (ITS1 and ITS2) differentially abundant genera in bulk soil (Sb) and rhizosphere (Sr) categorized by treatment (allium ’A’, banana ’B’, co- cultivation ’AB’ and control ’C’). Rarefied (Abundance), and Rarefied Relative (Rel-Ab) Abundances are reported.

| ITS | Soil type | Genus | A | | AB | | B | | C | |
|---|---|---|---|---|---|---|---|---|---|---|
| | | | **Abundance** | **Rel-Ab** | **Abundance** | **Rel-Ab** | **Abundance** | **Rel-Ab** | **Abundance** | **Rel-Ab** |
| **1** | **Sb** | Acremonium | 31.75 ± 26.75 | 25.2 ± 21.23 | 4.40 ± 10.92 | 3.49 ± 8.66 | 0.90 ± 1.29 | 0.71 ± 1.02 | 4.30 ± 7.80 | 3.41 ± 6.19 |
| | | Aspergillus | 1.00 ± 1.07 | 0.79 ± 0.85 | 0.10 ± 0.32 | 0.08 ± 0.25 | 0.10 ± 0.32 | 0.08 ± 0.25 | 0.20 ± 0.63 | 0.16 ± 0.50 |
| | **Sr** | Apodus | 2.00 ± 3.35 | 0.06 ± 0.11 | 1.33 ± 2.69 | 0.04 ± 0.09 | 4.89 ± 2.98 | 0.16 ± 0.10 | - | - |
| | | Ascobolus | 16.67 ± 29.61 | 0.54 ± 0.96 | 8.67 ± 10.72 | 0.28 ± 0.35 | 63.56 ± 76.72 | 2.06 ± 2.49 | - | - |
| | | Leptoxyphium | 2.44 ± 2.30 | 0.08 ± 0.07 | 0.00 ± 0.00 | 0.00 ± 0.00 | 0.00 ± 0.00 | 0.00 ± 0.00 | - | - |
| | | Mortierella | 150.67 ± 162.92 | 4.89 ± 5.28 | 8.56 ± 12.31 | 0.28 ± 0.40 | 11.44 ± 25.61 | 0.37 ± 0.83 | - | - |
| | | Tausonia | 4.33 ± 8.34 | 0.14 ± 0.27 | 3.33 ± 5.77 | 0.11 ± 0.19 | 21.67 ± 19.54 | 0.70 ± 0.63 | - | - |
| | | Trichoderma | 23.22 ± 19.06 | 0.75 ± 0.62 | 4.56 ± 5.61 | 0.15 ± 0.18 | 5.78 ± 10.94 | 0.19 ± 0.35 | - | - |
| **2** | **Sb** | Scedosporium | 22.50 ± 9.48 | 1.15 ± 0.48 | 60.44 ± 47.74 | 3.09 ± 2.44 | 18.75 ± 25.24 | 0.96 ± 1.29 | 10.22 ± 8.18 | 0.52 ± 0.42 |
| | | Wallrothiella | 0.17 ± 0.41 | 0.01 ± 0.02 | 2.44 ± 2.70 | 0.13 ± 0.14 | 0.00 ± 0.00 | 0.00 ± 0.00 | 0.11 ± 0.33 | 0.01 ± 0.02 |
| | **Sr** | Apiotrichum | 157.88 ± 110.07 | 3.38 ± 2.36 | 10.57 ± 21.60 | 0.23 ± 0.46 | 65.89 ± 147.58 | 1.41 ± 3.16 | - | - |
| | | Aquapteridospora | 8.12 ± 6.51 | 0.17 ± 0.14 | 2.00 ± 3.61 | 0.04 ± 0.08 | 0.89 ± 1.27 | 0.02 ± 0.03 | - | - |
| | | Arthrobotrys | 39.12 ± 30.14 | 0.84 ± 0.65 | 6.29 ± 4.89 | 0.13 ± 0.10 | 8.56 ± 10.42 | 0.18 ± 0.22 | - | - |
| | | Chloridium | 11.50 ± 13.47 | 0.25 ± 0.29 | 0.00 ± 0.00 | 0.00 ± 0.00 | 0.56 ± 1.67 | 0.01 ± 0.04 | - | - |
| | | Coprinopsis | 64.88 ± 35.51 | 1.39 ± 0.76 | 15.00 ± 26.21 | 0.32 ± 0.56 | 6.11 ± 12.67 | 0.13 ± 0.27 | - | - |
| | | Cylindrocladiella | 2.38 ± 4.14 | 0.05 ± 0.09 | 0.00 ± 0.00 | 0.00 ± 0.00 | 0.00 ± 0.00 | 0.00 ± 0.00 | - | - |
| | | Fusarium | 346.75 ± 248.84 | 7.43 ± 5.33 | 110.57 ± 197.14 | 2.37 ± 4.22 | 127.89 ± 255.45 | 2.74 ± 5.47 | - | - |
| | | Hyaloscypha | 3.38 ± 4.44 | 0.07 ± 0.10 | 0.00 ± 0.00 | 0.00 ± 0.00 | 0.00 ± 0.00 | 0.00 ± 0.00 | - | - |
| | | Lasiodiplodia | 11.00 ± 19.50 | 0.24 ± 0.42 | 0.00 ± 0.00 | 0.00 ± 0.00 | 0.11 ± 0.33 | 0.00 ± 0.01 | - | - |
| | | Microascus | 5.38 ± 5.50 | 0.12 ± 0.12 | 0.43 ± 1.13 | 0.01 ± 0.02 | 0.89 ± 1.83 | 0.02 ± 0.04 | - | - |
| | | Mortierella | 111.12 ± 53.47 | 2.38 ± 1.14 | 6.29 ± 12.22 | 0.13 ± 0.26 | 49.44 ± 97.06 | 1.06 ± 2.08 | - | - |
| | | Mycothermus | 118.88 ± 53.03 | 2.55 ± 1.14 | 39.43 ± 78.30 | 0.84 ± 1.68 | 17.67 ± 32.20 | 0.38 ± 0.69 | - | - |
| | | Scedosporium | 171.88 ± 149.85 | 3.68 ± 3.21 | 15.43 ± 31.43 | 0.33 ± 0.67 | 9.00 ± 16.03 | 0.19 ± 0.34 | - | - |
| | | Tausonia | 3.62 ± 6.46 | 0.08 ± 0.14 | 3.29 ± 3.55 | 0.07 ± 0.08 | 23.00 ± 25.93 | 0.49 ± 0.56 | - | - |
| | | Zopfiella | 10.88 ± 15.33 | 0.23 ± 0.33 | 32.57 ± 35.02 | 0.70 ± 0.75 | 5.78 ± 6.08 | 0.12 ± 0.13 | - | - |

**Table S7:** Summary of bacterial (16S rRNA) biomarkers and linear discriminant analysis (LDA) for bacterial amplicon sequence variants (ASVs) in bulk (Sb) and rhizosphere (Sr). ’Tx’ indicates the treatment in which the ASV exhibited significant abundance (allium ’A’, banana ’B’, co-cultivation ’AB’ and control ’C’). ’FDR’ stands for false discovery rate.

| Soil type | ASV | Taxonomy | | | | | | LDA upper | LDA mean | LDA lower | Tx | p - Value | FDR* |
|---|---|---|---|---|---|---|---|---|---|---|---|---|---|
| | | Phylum | Class | Order | Family | Genus | Species | | | | | | |
| Sb | ASV1 | Cyanobacteria | Cyanobacteriia | Geitlerinematales | Geitlerinemataceae | Geitlerinema | - | 4.121 | 4.055 | 3.977 | A | 0.000 | 0.000 |
| | ASV1009 | Proteobacteria | Gammaproteobacteria | Burkholderiales | Nitrosomonadaceae | - | - | 2.502 | 2.455 | 2.404 | AB | 0.001 | 0.041 |
| | ASV11 | Proteobacteria | Alphaproteobacteria | Rhizobiales | Rhizobiaceae | - | - | 3.572 | 3.521 | 3.464 | A | 0.000 | 0.016 |
| | ASV116 | Actinobacteriota | Actinobacteria | Micrococcales | Microbacteriaceae | Microbacterium | - | 3.094 | 3.035 | 2.968 | A | 0.000 | 0.005 |
| | ASV1231 | Planctomycetota | Planctomycetes | Planctomycetales | Schlesneriaceae | Planctopirus | - | 2.402 | 2.356 | 2.306 | AB | 0.000 | 0.017 |
| | ASV1268 | Bacteroidota | Bacteroidia | Chitinophagales | Chitinophagaceae | Flavihumibacter | - | 2.298 | 2.236 | 2.165 | A | 0.001 | 0.036 |
| | ASV143 | Actinobacteriota | Actinobacteria | Propionibacteriales | Nocardioidaceae | Marmoricola | - | 2.672 | 2.639 | 2.603 | C | 0.000 | 0.033 |
| | ASV1492 | Actinobacteriota | Acidimicrobiia | Actinomarinales | - | - | - | 2.255 | 2.214 | 2.169 | A | 0.001 | 0.047 |
| | ASV1573 | Actinobacteriota | Thermoleophilia | Gaiellales | - | - | - | 2.119 | 2.074 | 2.024 | A | 0.001 | 0.040 |
| | ASV1589 | Acidobacteriota | Blastocatellia | Blastocatellales | Blastocatellaceae | - | - | 2.262 | 2.201 | 2.131 | A | 0.001 | 0.036 |
| | ASV1842 | Proteobacteria | Alphaproteobacteria | Sphingomonadales | Sphingomonadaceae | Sphingomonas | - | 2.059 | 2.020 | 1.977 | C | 0.001 | 0.050 |
| | ASV1879 | Acidobacteriota | Vicinamibacteria | Vicinamibacterales | - | - | - | 2.090 | 2.036 | 1.974 | A | 0.000 | 0.016 |
| | ASV270 | Chloroflexi | Anaerolineae | Anaerolineales | Anaerolineaceae | - | - | 2.810 | 2.772 | 2.730 | AB | 0.001 | 0.047 |
| | ASV308 | Bacteroidota | Bacteroidia | Chitinophagales | Chitinophagaceae | Flavihumibacter | - | 2.777 | 2.722 | 2.659 | A | 0.000 | 0.016 |
| | ASV33 | Proteobacteria | Alphaproteobacteria | Sphingomonadales | | - | - | 3.137 | 3.101 | 3.061 | C | 0.000 | 0.017 |
| | ASV3352 | Proteobacteria | Alphaproteobacteria | Rhizobiales | Rhizobiaceae | - | - | 2.100 | 2.053 | 1.999 | C | 0.000 | 0.028 |
| | ASV344 | Proteobacteria | Alphaproteobacteria | Rhizobiales | Rhizobiaceae | - | - | 2.638 | 2.570 | 2.491 | A | 0.001 | 0.036 |
| | ASV415 | Proteobacteria | Alphaproteobacteria | Sphingomonadales | Sphingomonadaceae | Sphingobium | - | 2.754 | 2.713 | 2.667 | C | 0.000 | 0.012 |
| | ASV543 | Proteobacteria | Alphaproteobacteria | Caulobacterales | Caulobacteraceae | Brevundimonas | - | 2.606 | 2.562 | 2.513 | A | 0.001 | 0.036 |
| | ASV617 | Proteobacteria | Alphaproteobacteria | Rhizobiales | Devosiaceae | Devosia | - | 2.452 | 2.410 | 2.364 | A | 0.001 | 0.038 |
| | ASV675 | Bacteroidota | Bacteroidia | Chitinophagales | Chitinophagaceae | Flavihumibacter | - | 2.521 | 2.468 | 2.408 | A | 0.001 | 0.043 |
| | ASV725 | Actinobacteriota | Actinobacteria | Propionibacteriales | Nocardioidaceae | Nocardioides | - | 2.512 | 2.456 | 2.392 | A | 0.001 | 0.048 |
| | ASV747 | Acidobacteriota | Blastocatellia | Pyrinomonadales | Pyrinomonadaceae | - | - | 2.608 | 2.556 | 2.497 | AB | 0.000 | 0.015 |
| | ASV841 | Cyanobacteria | Cyanobacteriia | Phormidesmiales | Nodosilineaceae | - | - | 2.433 | 2.383 | 2.327 | A | 0.000 | 0.022 |
| | ASV980 | Proteobacteria | Gammaproteobacteria | Burkholderiales | Comamonadaceae | Rhizobacter | - | 2.481 | 2.417 | 2.342 | B | 0.000 | 0.028 |
| Sr | ASV103 | Acidobacteriota | Blastocatellia | - | - | - | - | 3.169 | 3.120 | 3.066 | A | 0.006 | 0.174 |
| | ASV1223 | Proteobacteria | Alphaproteobacteria | - | - | - | - | 3.107 | 3.039 | 2.959 | A | 0.009 | 0.200 |
| | ASV1331 | Actinobacteriota | Acidimicrobiia | Microtrichales | Ilumatobacteraceae | - | - | 3.142 | 3.073 | 2.991 | B | 0.005 | 0.172 |
| | ASV167 | Proteobacteria | Alphaproteobacteria | Rhizobiales | Xanthobacteraceae | - | - | 3.146 | 3.097 | 3.042 | A | 0.009 | 0.200 |
| | ASV17 | Actinobacteriota | Acidimicrobiia | Actinomarinales | - | - | - | 3.271 | 3.221 | 3.165 | A | 0.004 | 0.149 |
| | ASV197 | Proteobacteria | Gammaproteobacteria | Pseudomonadales | Pseudomonadaceae | Pseudomonas | umsongensis | 3.250 | 3.200 | 3.143 | B | 0.007 | 0.186 |
| | ASV198 | Proteobacteria | Gammaproteobacteria | - | - | - | - | 3.265 | 3.214 | 3.156 | A | 0.000 | 0.120 |
| | ASV206 | Gemmatimonadota | Gemmatimonadetes | Gemmatimonadales | Gemmatimonadaceae | - | - | 3.165 | 3.109 | 3.046 | A | 0.002 | 0.134 |
| | ASV208 | Acidobacteriota | Vicinamibacteria | Vicinamibacterales | Vicinamibacteraceae | - | - | 3.198 | 3.149 | 3.093 | A | 0.000 | 0.120 |
| | ASV249 | Proteobacteria | Gammaproteobacteria | Burkholderiales | Comamonadaceae | Ramlibacter | - | 3.174 | 3.118 | 3.053 | A | 0.005 | 0.172 |
| | ASV266 | Proteobacteria | Gammaproteobacteria | Burkholderiales | Comamonadaceae | Aquabacterium | - | 3.267 | 3.213 | 3.153 | B | 0.000 | 0.104 |
| | ASV304 | Proteobacteria | Alphaproteobacteria | Rhizobiales | - | - | - | 3.176 | 3.091 | 2.985 | A | 0.009 | 0.200 |
| | ASV329 | Actinobacteriota | Thermoleophilia | Gaiellales | - | - | - | 3.216 | 3.154 | 3.081 | A | 0.003 | 0.141 |
| | ASV339 | Patescibacteria | Saccharimonadia | Saccharimonadales | Saccharimonadales | - | - | 3.219 | 3.169 | 3.113 | B | 0.001 | 0.120 |
| | ASV407 | Proteobacteria | Gammaproteobacteria | Burkholderiales | - | - | - | 3.073 | 3.002 | 2.917 | A | 0.008 | 0.197 |
| | ASV41 | Chloroflexi | - | - | - | - | - | 3.259 | 3.216 | 3.168 | A | 0.002 | 0.137 |
| | ASV43 | Chloroflexi | - | - | - | - | - | 3.199 | 3.149 | 3.093 | A | 0.007 | 0.186 |
| | ASV45 | Proteobacteria | Alphaproteobacteria | Sphingomonadales | Sphingomonadaceae | Sphingomonas | - | 3.207 | 3.156 | 3.098 | A | 0.001 | 0.120 |

| | | | | | | | | | | | | | |
|---|---|---|---|---|---|---|---|---|---|---|---|---|---|
| | ASV469 | Acidobacteriota | Acidobacteriae | - | - | - | - | 3.060 | 2.995 | 2.918 | A | 0.003 | 0.141 |
| | ASV478 | Chloroflexi | Dehalococcoidia | - | - | - | - | 3.127 | 3.068 | 2.999 | A | 0.002 | 0.134 |
| | ASV48 | Actinobacteriota | Acidimicrobiia | Actinomarinales | - | - | - | 3.212 | 3.156 | 3.093 | A | 0.005 | 0.172 |
| | ASV483 | Proteobacteria | Gammaproteobacteria | - | - | - | - | 3.175 | 3.111 | 3.037 | A | 0.001 | 0.120 |
| | ASV504 | Actinobacteriota | Acidimicrobiia | - | - | - | - | 3.007 | 2.943 | 2.869 | A | 0.009 | 0.200 |
| | ASV56 | Proteobacteria | Gammaproteobacteria | - | - | - | - | 3.133 | 3.085 | 3.031 | A | 0.003 | 0.147 |
| | ASV568 | Planctomycetota | Planctomycetes | Pirellulales | Pirellulaceae | - | - | 3.102 | 3.045 | 2.978 | A | 0.003 | 0.141 |
| | ASV620 | Actinobacteriota | Actinobacteria | Micrococcales | Micrococcaceae | Pseudarthrobacter | - | 3.089 | 3.036 | 2.976 | B | 0.004 | 0.150 |
| | ASV652 | Acidobacteriota | Vicinamibacteria | - | - | - | - | 2.972 | 2.905 | 2.824 | A | 0.009 | 0.200 |
| | ASV671 | Patescibacteria | - | - | - | - | - | 3.075 | 3.018 | 2.952 | A | 0.002 | 0.138 |
| | ASV7 | Proteobacteria | Alphaproteobacteria | Rhizobiales | Xanthobacteraceae | Pseudolabrys | - | 3.488 | 3.438 | 3.381 | A | 0.009 | 0.200 |
| | ASV75 | Chloroflexi | - | - | - | - | - | 3.065 | 3.012 | 2.952 | A | 0.004 | 0.150 |
| | ASV793 | Actinobacteriota | Actinobacteria | Micrococcales | Micrococcaceae | - | - | 3.016 | 2.958 | 2.892 | B | 0.005 | 0.172 |
| | ASV804 | Actinobacteriota | Acidimicrobiia | Microtrichales | Ilumatobacteraceae | Ilumatobacter | - | 3.090 | 3.026 | 2.952 | A | 0.009 | 0.200 |
| | ASV874 | Acidobacteriota | Vicinamibacteria | Vicinamibacterales | - | - | - | 3.142 | 3.088 | 3.027 | B | 0.003 | 0.147 |
| | ASV895 | Proteobacteria | Gammaproteobacteria | Burkholderiales | Oxalobacteraceae | Noviherbaspirillum | - | 3.161 | 3.098 | 3.023 | B | 0.004 | 0.150 |
| | ASV97 | Actinobacteriota | Acidimicrobiia | Actinomarinales | - | - | - | 3.131 | 3.086 | 3.035 | A | 0.003 | 0.141 |

**Table S8:** Abundance of bacterial (16S rRNA) biomarkers in bulk soil (Sb) and rhizosphere (Sr) categorized by treatment (allium ’A’, banana ’B’, co- cultivation ’AB’ and control ’C’). Rarefied (Abundance), and Rarefied Relative (Rel-Ab) Abundances are reported.

| Soil type | ASV | A | | AB | | B | | C | |
|---|---|---|---|---|---|---|---|---|---|
| | | Abundance | Rel-Ab | Abundance | Rel-Ab | Abundance | Rel-Ab | Abundance | Rel-Ab |
| **Sb** | ASV1 | 258.44 ± 447.54 | 4.41 ± 7.63 | 1.00 ± 2.83 | 0.02 ± 0.05 | 0.00 ± 0.00 | 0.00 ± 0.00 | 0.00 ± 0.00 | 0.00 ± 0.00 |
| | ASV1009 | 0.11 ± 0.33 | 0.00 ± 0.01 | 5.50 ± 3.59 | 0.09 ± 0.06 | 1.30 ± 1.95 | 0.02 ± 0.03 | 0.00 ± 0.00 | 0.00 ± 0.00 |
| | ASV11 | 73.56 ± 84.53 | 1.25 ± 1.44 | 7.88 ± 21.48 | 0.13 ± 0.37 | 13.80 ± 26.65 | 0.24 ± 0.45 | 0.00 ± 0.00 | 0.00 ± 0.00 |
| | ASV116 | 24.89 ± 21.13 | 0.42 ± 0.36 | 0.50 ± 1.41 | 0.01 ± 0.02 | 0.20 ± 0.63 | 0.00 ± 0.01 | 1.20 ± 2.30 | 0.02 ± 0.04 |
| | ASV1231 | 0.11 ± 0.33 | 0.00 ± 0.01 | 4.75 ± 2.82 | 0.08 ± 0.05 | 1.20 ± 1.62 | 0.02 ± 0.03 | 0.00 ± 0.00 | 0.00 ± 0.00 |
| | ASV1268 | 3.78 ± 4.82 | 0.06 ± 0.08 | 0.00 ± 0.00 | 0.00 ± 0.00 | 0.00 ± 0.00 | 0.00 ± 0.00 | 0.00 ± 0.00 | 0.00 ± 0.00 |
| | ASV143 | 6.44 ± 4.69 | 0.11 ± 0.08 | 5.50 ± 3.46 | 0.09 ± 0.06 | 10.10 ± 3.18 | 0.17 ± 0.05 | 14.00 ± 3.43 | 0.24 ± 0.06 |
| | ASV1492 | 3.00 ± 2.4 | 0.05 ± 0.04 | 0.00 ± 0.00 | 0.00 ± 0.00 | 0.50 ± 0.85 | 0.01 ± 0.01 | 0.70 ± 1.49 | 0.01 ± 0.03 |
| | ASV1573 | 1.78 ± 2.05 | 0.03 ± 0.03 | 0.00 ± 0.00 | 0.00 ± 0.00 | 0.00 ± 0.00 | 0.00 ± 0.00 | 0.30 ± 0.95 | 0.01 ± 0.02 |
| | ASV1589 | 3.44 ± 4.48 | 0.06 ± 0.08 | 0.00 ± 0.00 | 0.00 ± 0.00 | 0.00 ± 0.00 | 0.00 ± 0.00 | 0.00 ± 0.00 | 0.00 ± 0.00 |
| | ASV1842 | 0.00 ± 0.00 | 0.00 ± 0.00 | 0.00 ± 0.00 | 0.00 ± 0.00 | 0.40 ± 0.84 | 0.01 ± 0.01 | 1.70 ± 1.83 | 0.03 ± 0.03 |
| | ASV1879 | 2.00 ± 2.06 | 0.03 ± 0.04 | 0.00 ± 0.00 | 0.00 ± 0.00 | 0.00 ± 0.00 | 0.00 ± 0.00 | 0.00 ± 0.00 | 0.00 ± 0.00 |
| | ASV270 | 1.56 ± 2.65 | 0.03 ± 0.05 | 13.50 ± 7.56 | 0.23 ± 0.13 | 6.60 ± 4.25 | 0.11 ± 0.07 | 2.50 ± 3.81 | 0.04 ± 0.06 |
| | ASV308 | 11.78 ± 12.4 | 0.20 ± 0.21 | 0.25 ± 0.71 | 0.00 ± 0.01 | 1.50 ± 4.74 | 0.03 ± 0.08 | 0.10 ± 0.32 | 0.00 ± 0.01 |
| | ASV33 | 26.44 ± 9.41 | 0.45 ± 0.16 | 15.62 ± 12.25 | 0.27 ± 0.21 | 18.4 ± 8.09 | 0.31 ± 0.14 | 41.30 ± 8.43 | 0.70 ± 0.14 |
| | ASV3352 | 0.00 ± 0.00 | 0.00 ± 0.00 | 0.00 ± 0.00 | 0.00 ± 0.00 | 0.00 ± 0.00 | 0.00 ± 0.00 | 1.80 ± 2.74 | 0.03 ± 0.05 |
| | ASV344 | 8.33 ± 15.94 | 0.14 ± 0.27 | 0.00 ± 0.00 | 0.00 ± 0.00 | 0.00 ± 0.00 | 0.00 ± 0.00 | 0.00 ± 0.00 | 0.00 ± 0.00 |
| | ASV415 | 0.00 ± 0.00 | 0.00 ± 0.00 | 0.00 ± 0.00 | 0.00 ± 0.00 | 3.50 ± 4.03 | 0.06 ± 0.07 | 10.8 ± 6.36 | 0.18 ± 0.11 |
| | ASV543 | 7.56 ± 6.54 | 0.13 ± 0.11 | 0.50 ± 1.41 | 0.01 ± 0.02 | 0.00 ± 0.00 | 0.00 ± 0.00 | 2.50 ± 7.91 | 0.04 ± 0.13 |
| | ASV617 | 4.89 ± 4.04 | 0.08 ± 0.07 | 2.25 ± 6.36 | 0.04 ± 0.11 | 0.00 ± 0.00 | 0.00 ± 0.00 | 0.50 ± 1.58 | 0.01 ± 0.03 |
| | ASV675 | 6.33 ± 8.87 | 0.11 ± 0.15 | 0.25 ± 0.71 | 0.00 ± 0.01 | 0.00 ± 0.00 | 0.00 ± 0.00 | 1.10 ± 2.42 | 0.02 ± 0.04 |
| | ASV725 | 6.22 ± 8.41 | 0.11 ± 0.14 | 0.00 ± 0.00 | 0.00 ± 0.00 | 0.30 ± 0.95 | 0.01 ± 0.02 | 0.70 ± 1.49 | 0.01 ± 0.03 |
| | ASV747 | 0.00 ± 0.00 | 0.00 ± 0.00 | 7.75 ± 5.95 | 0.13 ± 0.10 | 1.20 ± 2.10 | 0.02 ± 0.04 | 0.00 ± 0.00 | 0.00 ± 0.00 |
| | ASV841 | 5.22 ± 4.32 | 0.09 ± 0.07 | 0.00 ± 0.00 | 0.00 ± 0.00 | 1.10 ± 2.60 | 0.02 ± 0.04 | 0.00 ± 0.00 | 0.00 ± 0.00 |
| | ASV980 | 0.00 ± 0.00 | 0.00 ± 0.00 | 0.00 ± 0.00 | 0.00 ± 0.00 | 6.50 ± 9.12 | 0.11 ± 0.16 | 0.00 ± 0.00 | 0.00 ± 0.00 |
| **Sr** | ASV103 | 2.89 ± 2.09 | 0.32 ± 0.23 | 0.71 ± 1.25 | 0.08 ± 0.14 | 0.43 ± 0.79 | 0.05 ± 0.09 | - | - |
| | ASV1223 | 0.67 ± 0.71 | 0.07 ± 0.08 | 0.00 ± 0.00 | 0.00 ± 0.00 | 0.00 ± 0.00 | 0.00 ± 0.00 | - | - |
| | ASV1331 | 0.00 ± 0.00 | 0.00 ± 0.00 | 0.00 ± 0.00 | 0.00 ± 0.00 | 1.00 ± 1.41 | 0.11 ± 0.16 | - | - |
| | ASV167 | 2.89 ± 1.83 | 0.32 ± 0.20 | 0.71 ± 1.89 | 0.08 ± 0.21 | 0.43 ± 0.79 | 0.05 ± 0.09 | - | - |
| | ASV17 | 5.56 ± 3.09 | 0.61 ± 0.34 | 1.57 ± 1.51 | 0.17 ± 0.17 | 2.00 ± 1.29 | 0.22 ± 0.14 | - | - |
| | ASV197 | 0.56 ± 1.13 | 0.06 ± 0.12 | 0.57 ± 1.13 | 0.06 ± 0.12 | 3.71 ± 2.29 | 0.41 ± 0.25 | - | - |
| | ASV198 | 2.44 ± 1.59 | 0.27 ± 0.17 | 0.29 ± 0.76 | 0.03 ± 0.08 | 0.00 ± 0.00 | 0.00 ± 0.00 | - | - |
| | ASV206 | 2.44 ± 1.81 | 0.27 ± 0.20 | 0.57 ± 1.13 | 0.06 ± 0.12 | 0.00 ± 0.00 | 0.00 ± 0.00 | - | - |
| | ASV208 | 2.56 ± 1.59 | 0.28 ± 0.17 | 0.43 ± 0.79 | 0.05 ± 0.09 | 0.00 ± 0.00 | 0.00 ± 0.00 | - | - |
| | ASV249 | 1.11 ± 0.78 | 0.12 ± 0.09 | 0.29 ± 0.49 | 0.03 ± 0.05 | 0.00 ± 0.00 | 0.00 ± 0.00 | - | - |
| | ASV266 | 0.00 ± 0.00 | 0.00 ± 0.00 | 0.14 ± 0.38 | 0.02 ± 0.04 | 3.71 ± 2.29 | 0.41 ± 0.25 | - | - |
| | ASV304 | 0.78 ± 0.97 | 0.09 ± 0.11 | 0.00 ± 0.00 | 0.00 ± 0.00 | 0.00 ± 0.00 | 0.00 ± 0.00 | - | - |
| | ASV329 | 0.89 ± 0.78 | 0.10 ± 0.09 | 0.00 ± 0.00 | 0.00 ± 0.00 | 0.00 ± 0.00 | 0.00 ± 0.00 | - | - |
| | ASV339 | 0.22 ± 0.44 | 0.02 ± 0.05 | 0.57 ± 0.79 | 0.06 ± 0.09 | 3.00 ± 1.41 | 0.33 ± 0.16 | - | - |
| | ASV407 | 1.11 ± 0.93 | 0.12 ± 0.10 | 0.14 ± 0.38 | 0.02 ± 0.04 | 0.00 ± 0.00 | 0.00 ± 0.00 | - | - |

| | | | | | | | | | |
|---|---|---|---|---|---|---|---|---|---|
| | ASV41 | 4.56 ± 1.33 | 0.50 ± 0.15 | 1.71 ± 2.21 | 0.19 ± 0.24 | 1.00 ± 0.58 | 0.11 ± 0.06 | - | - |
| | ASV43 | 3.56 ± 2.40 | 0.39 ± 0.26 | 0.86 ± 0.69 | 0.09 ± 0.08 | 1.29 ± 1.38 | 0.14 ± 0.15 | - | - |
| | ASV45 | 2.89 ± 1.90 | 0.32 ± 0.21 | 0.57 ± 0.53 | 0.06 ± 0.06 | 0.29 ± 0.49 | 0.03 ± 0.05 | - | - |
| | ASV469 | 1.00 ± 1.00 | 0.11 ± 0.11 | 0.00 ± 0.00 | 0.00 ± 0.00 | 0.00 ± 0.00 | 0.00 ± 0.00 | - | - |
| | ASV478 | 2.22 ± 1.72 | 0.24 ± 0.19 | 0.14 ± 0.38 | 0.02 ± 0.04 | 0.00 ± 0.00 | 0.00 ± 0.00 | - | - |
| | ASV48 | 3.00 ± 1.73 | 0.33 ± 0.19 | 0.71 ± 1.25 | 0.08 ± 0.14 | 0.29 ± 0.49 | 0.03 ± 0.05 | - | - |
| | ASV483 | 0.89 ± 0.60 | 0.10 ± 0.07 | 0.00 ± 0.00 | 0.00 ± 0.00 | 0.00 ± 0.00 | 0.00 ± 0.00 | - | - |
| | ASV504 | 1.11 ± 1.17 | 0.12 ± 0.13 | 0.00 ± 0.00 | 0.00 ± 0.00 | 0.00 ± 0.00 | 0.00 ± 0.00 | - | - |
| | ASV56 | 2.89 ± 1.54 | 0.32 ± 0.17 | 0.43 ± 0.79 | 0.05 ± 0.09 | 1.29 ± 0.76 | 0.14 ± 0.08 | - | - |
| | ASV568 | 1.22 ± 1.20 | 0.13 ± 0.13 | 0.00 ± 0.00 | 0.00 ± 0.00 | 0.00 ± 0.00 | 0.00 ± 0.00 | - | - |
| | ASV620 | 0.33 ± 0.71 | 0.04 ± 0.08 | 0.00 ± 0.00 | 0.00 ± 0.00 | 1.29 ± 0.95 | 0.14 ± 0.10 | - | - |
| | ASV652 | 1.00 ± 1.22 | 0.11 ± 0.13 | 0.00 ± 0.00 | 0.00 ± 0.00 | 0.00 ± 0.00 | 0.00 ± 0.00 | - | - |
| | ASV671 | 1.22 ± 0.83 | 0.13 ± 0.09 | 0.14 ± 0.38 | 0.02 ± 0.04 | 0.00 ± 0.00 | 0.00 ± 0.00 | - | - |
| | ASV7 | 10.78 ± 4.63 | 1.18 ± 0.51 | 4.00 ± 2.83 | 0.44 ± 0.31 | 4.43 ± 2.82 | 0.49 ± 0.31 | - | - |
| | ASV75 | 2.00 ± 1.58 | 0.22 ± 0.17 | 0.43 ± 0.53 | 0.05 ± 0.06 | 0.29 ± 0.49 | 0.03 ± 0.05 | - | - |
| | ASV793 | 0.00 ± 0.00 | 0.00 ± 0.00 | 0.00 ± 0.00 | 0.00 ± 0.00 | 1.00 ± 1.00 | 0.11 ± 0.11 | - | - |
| | ASV804 | 0.78 ± 0.83 | 0.09 ± 0.09 | 0.00 ± 0.00 | 0.00 ± 0.00 | 0.00 ± 0.00 | 0.00 ± 0.00 | - | - |
| | ASV874 | 0.00 ± 0.00 | 0.00 ± 0.00 | 0.14 ± 0.38 | 0.02 ± 0.04 | 1.43 ± 1.13 | 0.16 ± 0.12 | - | - |
| | ASV895 | 0.00 ± 0.00 | 0.00 ± 0.00 | 0.14 ± 0.38 | 0.02 ± 0.04 | 1.14 ± 0.90 | 0.13 ± 0.10 | - | - |
| | ASV97 | 2.78 ± 1.64 | 0.31 ± 0.18 | 0.86 ± 0.90 | 0.09 ± 0.10 | 0.29 ± 0.76 | 0.03 ± 0.08 | - | - |

**Table S9:** Summary of fungal (ITS1 and ITS2) biomarkers and linear discriminant analysis (LDA) for fungal amplicon sequence variants (ASVs) in bulk (Sb) and rhizosphere (Sr). 'Tx' indicates the treatment in which the ASV exhibited significant abundance. 'FDR' stands for false discovery rate.

| ITS | Soil type | ASV | Taxonomy | | | | | | LDA upper | LDA mean | LDA lower | Plant | p - Value | FDR |
|---|---|---|---|---|---|---|---|---|---|---|---|---|---|---|
| | | | Phylum | Class | Order | Family | Genus | Species | | | | | | |
| **1** | Sb | ASV417 | Ascomycota | Sordariomycetes | Hypocreales | Incertae sedis | Acremonium | exuviarum | 3.439 | 3.377 | 3.306 | A | 0.008 | 0.532 |
| | | ASV46 | Ascomycota | Sordariomycetes | Hypocreales | Nectriaceae | - | - | 3.593 | 3.544 | 3.489 | A | 0.014 | 0.596 |
| | Sr | ASV132 | Ascomycota | Sordariomycetes | Microascales | - | - | - | 2.968 | 2.914 | 2.853 | A | 0.008 | 0.329 |
| | | ASV153 | Ascomycota | Sordariomycetes | Hypocreales | Hypocreaceae | Trichoderma | erinaceum | 2.952 | 2.884 | 2.805 | A | 0.024 | 0.402 |
| | | ASV16 | Ascomycota | Sordariomycetes | Sordariales | Chaetomiaceae | Mycothermus | thermophilus | 3.844 | 3.798 | 3.747 | A | 0.004 | 0.313 |
| | | ASV161 | Ascomycota | Sordariomycetes | Microascales | Microascaceae | Cephalotrichum | stemonitis | 3.045 | 2.986 | 2.919 | B | 0.034 | 0.447 |
| | | ASV189 | Ascomycota | - | - | - | - | - | 2.710 | 2.647 | 2.573 | A | 0.003 | 0.313 |
| | | ASV2 | Ascomycota | Sordariomycetes | Hypocreales | Incertae sedis | Acremonium | exuviarum | 3.551 | 3.470 | 3.372 | A | 0.011 | 0.344 |
| | | ASV210 | Ascomycota | Sordariomycetes | Glomerellales | Plectosphaerellaceae | Plectosphaerella | cucumerina | 2.983 | 2.920 | 2.847 | B | 0.011 | 0.344 |
| | | ASV211 | Ascomycota | Sordariomycetes | Sordariales | Lasiosphaeriaceae | Apodus | deciduus | 2.638 | 2.591 | 2.538 | B | 0.009 | 0.344 |
| | | ASV213 | Mortierellomycota | Mortierellomycetes | Mortierellales | Mortierellaceae | Mortierella | - | 2.643 | 2.575 | 2.494 | A | 0.011 | 0.344 |
| | | ASV216 | Ascomycota | Sordariomycetes | Microascales | Microascaceae | Cephalotrichum | stemonitis | 2.974 | 2.909 | 2.833 | B | 0.011 | 0.344 |
| | | ASV466 | Basidiomycota | Tremellomycetes | Filobasidiales | Piskurozymaceae | Solicoccozyma | terricola | 2.652 | 2.570 | 2.468 | B | 0.011 | 0.344 |
| | | ASV50 | Basidiomycota | Agaricomycetes | Trechisporales | Hydnodontaceae | Subulicystidium | brachysporum | 3.361 | 3.302 | 3.234 | B | 0.033 | 0.447 |
| | | ASV66 | Mortierellomycota | Mortierellomycetes | Mortierellales | Mortierellaceae | Mortierella | exigua | 3.655 | 3.559 | 3.435 | A | 0.011 | 0.344 |
| | | ASV69 | Basidiomycota | Tremellomycetes | Cystofilobasidiales | Mrakiaceae | Tausonia | pullulans | 3.366 | 3.308 | 3.240 | B | 0.004 | 0.313 |
| | | ASV715 | Aphelidiomycota | Aphelidiomycetes | - | - | - | - | 2.694 | 2.623 | 2.539 | B | 0.011 | 0.344 |
| | | ASV82 | Basidiomycota | Agaricomycetes | Trechisporales | Hydnodontaceae | Subulicystidium | brachysporum | 3.027 | 2.975 | 2.915 | B | 0.018 | 0.386 |
| **2** | Sb | ASV286 | Ascomycota | Sordariomycetes | Hypocreales | Incertae sedis | Acremonium | exuviarum | 3.251 | 3.207 | 3.158 | A | 0.008 | 0.204 |
| | Sr | ASV100 | Mortierellomycota | Mortierellomycetes | Mortierellales | Mortierellaceae | Mortierella | ambigua | 3.449 | 3.392 | 3.326 | A | 0.007 | 0.135 |
| | | ASV1012 | Basidiomycota | Agaricomycetes | - | - | - | - | 2.601 | 2.533 | 2.451 | A | 0.011 | 0.135 |
| | | ASV105 | Basidiomycota | Tremellomycetes | Cystofilobasidiales | Mrakiaceae | Tausonia | pullulans | 3.269 | 3.210 | 3.142 | B | 0.030 | 0.205 |
| | | ASV151 | Basidiomycota | Agaricomycetes | Agaricales | Psathyrellaceae | Coprinopsis | annulopora | 3.113 | 3.052 | 2.982 | A | 0.007 | 0.135 |
| | | ASV165 | Ascomycota | Sordariomycetes | Incertae sedis | Incertae sedis | Aquapteridospora | bambusinum | 2.865 | 2.801 | 2.727 | A | 0.006 | 0.135 |
| | | ASV178 | Ascomycota | Sordariomycetes | Microascales | Microascaceae | Scedosporium | prolificans | 3.294 | 3.230 | 3.156 | A | 0.013 | 0.154 |
| | | ASV183 | Basidiomycota | Agaricomycetes | Sebacinales | Serendipitaceae | Serendipita | - | 2.891 | 2.828 | 2.755 | AB | 0.004 | 0.135 |
| | | ASV218 | Ascomycota | Sordariomycetes | Sordariales | - | - | - | 3.414 | 3.343 | 3.258 | A | 0.021 | 0.188 |
| | | ASV233 | Ascomycota | Sordariomycetes | Microascales | Microascaceae | Scedosporium | dehoogii | 3.225 | 3.165 | 3.096 | A | 0.003 | 0.135 |
| | | ASV24 | Ascomycota | Sordariomycetes | Hypocreales | Nectriaceae | Fusarium | - | 3.824 | 3.771 | 3.709 | A | 0.022 | 0.188 |
| | | ASV26 | Basidiomycota | Tremellomycetes | Trichosporonales | Trichosporonaceae | Apiotrichum | dehoogii | 4.065 | 4.014 | 3.955 | A | 0.007 | 0.135 |
| | | ASV269 | Ascomycota | Sordariomycetes | Microascales | Halosphaeriaceae | - | - | 2.774 | 2.712 | 2.640 | A | 0.005 | 0.135 |
| | | ASV270 | Ascomycota | Dothideomycetes | Botryosphaeriales | Botryosphaeriaceae | Lasiodiplodia | gonubiensis | 2.976 | 2.903 | 2.815 | A | 0.008 | 0.135 |
| | | ASV28 | Ascomycota | Sordariomycetes | Sordariales | Chaetomiaceae | Mycothermus | thermophilus | 3.920 | 3.871 | 3.817 | A | 0.006 | 0.135 |
| | | ASV285 | Ascomycota | Pezizomycetes | Pezizales | Pyronemataceae | Lasiobolidium | orbiculoides | 2.996 | 2.925 | 2.841 | A | 0.011 | 0.135 |
| | | ASV333 | Ascomycota | Sordariomycetes | Microascales | Microascaceae | Enterocarpus | grenotii | 2.989 | 2.928 | 2.856 | A | 0.008 | 0.135 |
| | | ASV382 | Ascomycota | Sordariomycetes | Chaetosphaeriales | Chaetosphaeriaceae | Chloridium | submersum | 2.977 | 2.909 | 2.828 | A | 0.010 | 0.135 |
| | | ASV472 | Ascomycota | Sordariomycetes | Hypocreales | Nectriaceae | Cylindrocladiella | variabilis | 2.437 | 2.367 | 2.284 | A | 0.011 | 0.135 |
| | | ASV492 | Ascomycota | Orbiliomycetes | Orbiliales | Orbiliaceae | Arthrobotrys | - | 2.717 | 2.659 | 2.593 | A | 0.026 | 0.194 |
| | | ASV52 | Ascomycota | Dothideomycetes | Capnodiales | Cladosporiaceae | Cladosporium | cladosporioides | 2.987 | 2.922 | 2.847 | A | 0.003 | 0.135 |
| | | ASV53 | Ascomycota | Sordariomycetes | Hypocreales | Nectriaceae | Fusarium | solani | 3.555 | 3.497 | 3.431 | A | 0.016 | 0.178 |

| | | | | | | | | | | | | | | |
|---|---|---|---|---|---|---|---|---|---|---|---|---|---|---|
| | | ASV557 | Ascomycota | Sordariomycetes | Microascales | Microascaceae | Microascus | brevicaulis | 2.627 | 2.552 | 2.460 | A | 0.011 | 0.135 |
| | | ASV58 | Basidiomycota | Agaricomycetes | Agaricales | Psathyrellaceae | Coprinopsis | annulopora | 3.550 | 3.500 | 3.443 | A | 0.006 | 0.135 |
| | | ASV601 | Basidiomycota | Agaricomycetes | Russulales | - | - | - | 2.786 | 2.720 | 2.644 | A | 0.008 | 0.135 |
| | | ASV607 | Ascomycota | Sordariomycetes | Sordariales | Chaetomiaceae | Humicola | nigrescens | 3.078 | 2.988 | 2.874 | B | 0.007 | 0.135 |
| | | ASV616 | Basidiomycota | Agaricomycetes | Sebacinales | - | - | - | 2.750 | 2.676 | 2.587 | A | 0.007 | 0.135 |
| | | ASV70 | Ascomycota | Sordariomycetes | Microascales | Microascaceae | Scedosporium | - | 3.696 | 3.636 | 3.566 | A | 0.001 | 0.135 |
| | | ASV74 | Ascomycota | Eurotiomycetes | Eurotiales | - | - | - | 3.406 | 3.357 | 3.302 | A | 0.009 | 0.135 |
| | | ASV745 | Rozellomycota | - | - | - | - | - | 2.593 | 2.528 | 2.450 | A | 0.002 | 0.135 |
| | | ASV839 | Ascomycota | Leotiomycetes | Helotiales | Hyaloscyphaceae | Hyaloscypha | - | 2.558 | 2.486 | 2.399 | A | 0.010 | 0.135 |
| | | ASV99 | Ascomycota | Sordariomycetes | Hypocreales | Nectriaceae | Fusarium | - | 3.332 | 3.282 | 3.227 | A | 0.007 | 0.135 |

**Table S10:** Abundance of fungal (ITS1 and ITS2) biomarkers in bulk soil (Sb) and rhizosphere (Sr) categorized by treatment (allium 'A', banana 'B', co-cultivation 'AB' and control 'C'). Rarefied (Abundance) and Rarefied Relative (Rel-Ab) Abundances are reported.

| ITS | Soil type | ASV | Allium | | Allium + Banana | | Banana | | None | |
|---|---|---|---|---|---|---|---|---|---|---|
| | | | Abundance | Rel-Ab | Abundance | Rel-Ab | Abundance | Rel-Ab | Abundance | Rel-Ab |
| **1** | Sb | ASV417 | 0.38 ± 0.52 | 0.30 ± 0.41 | 0.00 ± 0.00 | 0.00 ± 0.00 | 0.00 ± 0.00 | 0.00 ± 0.00 | 0.00 ± 0.00 | 0.00 ± 0.00 |
| | | ASV46 | 1.75 ± 1.28 | 1.39 ± 1.02 | 0.60 ± 0.70 | 0.48 ± 0.55 | 0.30 ± 0.67 | 0.24 ± 0.54 | 0.40 ± 0.52 | 0.32 ± 0.41 |
| | Sr | ASV132 | 7.56 ± 8.88 | 0.24 ± 0.29 | 2.00 ± 4.92 | 0.06 ± 0.16 | 0.22 ± 0.67 | 0.01 ± 0.02 | - | - |
| | | ASV153 | 6.11 ± 9.69 | 0.20 ± 0.31 | 0.11 ± 0.33 | 0.00 ± 0.01 | 0.11 ± 0.33 | 0.00 ± 0.01 | - | - |
| | | ASV16 | 63.33 ± 50.49 | 2.05 ± 1.64 | 18.56 ± 26.95 | 0.60 ± 0.87 | 1.78 ± 2.99 | 0.06 ± 0.10 | - | - |
| | | ASV161 | 0.67 ± 2.00 | 0.02 ± 0.06 | 0.44 ± 1.33 | 0.01 ± 0.04 | 8.56 ± 11.47 | 0.28 ± 0.37 | - | - |
| | | ASV189 | 2.78 ± 3.70 | 0.09 ± 0.12 | 0.00 ± 0.00 | 0.00 ± 0.00 | 0.00 ± 0.00 | 0.00 ± 0.00 | - | - |
| | | ASV2 | 28.67 ± 62.16 | 0.93 ± 2.02 | 0.00 ± 0.00 | 0.00 ± 0.00 | 0.00 ± 0.00 | 0.00 ± 0.00 | - | - |
| | | ASV210 | 0.00 ± 0.00 | 0.00 ± 0.00 | 0.00 ± 0.00 | 0.00 ± 0.00 | 5.89 ± 14.01 | 0.19 ± 0.45 | - | - |
| | | ASV211 | 0.89 ± 1.45 | 0.03 ± 0.05 | 1.22 ± 2.39 | 0.04 ± 0.08 | 3.56 ± 2.30 | 0.12 ± 0.07 | - | - |
| | | ASV213 | 2.89 ± 5.28 | 0.09 ± 0.17 | 0.00 ± 0.00 | 0.00 ± 0.00 | 0.00 ± 0.00 | 0.00 ± 0.00 | - | - |
| | | ASV216 | 0.22 ± 0.67 | 0.01 ± 0.02 | 0.22 ± 0.44 | 0.01 ± 0.01 | 6.11 ± 8.61 | 0.20 ± 0.28 | - | - |
| | | ASV466 | 0.00 ± 0.00 | 0.00 ± 0.00 | 0.00 ± 0.00 | 0.00 ± 0.00 | 2.33 ± 3.54 | 0.08 ± 0.11 | - | - |
| | | ASV50 | 4.33 ± 6.63 | 0.14 ± 0.22 | 5.44 ± 8.66 | 0.18 ± 0.28 | 21.00 ± 24.39 | 0.68 ± 0.79 | - | - |
| | | ASV66 | 40.22 ± 105.13 | 1.30 ± 3.41 | 0.33 ± 1.00 | 0.01 ± 0.03 | 0.00 ± 0.00 | 0.00 ± 0.00 | - | - |
| | | ASV69 | 4.33 ± 8.34 | 0.14 ± 0.27 | 3.22 ± 5.47 | 0.10 ± 0.18 | 21.44 ± 19.08 | 0.70 ± 0.62 | - | - |
| | | ASV715 | 0.00 ± 0.00 | 0.00 ± 0.00 | 0.00 ± 0.00 | 0.00 ± 0.00 | 0.44 ± 0.53 | 0.01 ± 0.02 | - | - |
| | | ASV82 | 1.67 ± 5.00 | 0.05 ± 0.16 | 2.00 ± 3.04 | 0.06 ± 0.10 | 8.89 ± 8.24 | 0.29 ± 0.27 | - | - |
| **2** | Sb | ASV286 | 10.83 ± 8.35 | 0.55 ± 0.43 | 1.89 ± 2.89 | 0.10 ± 0.15 | 0.12 ± 0.35 | 0.01 ± 0.02 | 2.00 ± 6.00 | 0.10 ± 0.31 |
| | Sr | ASV100 | 38.25 ± 34.87 | 0.82 ± 0.75 | 2.86 ± 6.69 | 0.06 ± 0.14 | 9.33 ± 15.58 | 0.20 ± 0.33 | - | - |
| | | ASV1012 | 4.12 ± 5.72 | 0.09 ± 0.12 | 0.00 ± 0.00 | 0.00 ± 0.00 | 0.00 ± 0.00 | 0.00 ± 0.00 | - | - |
| | | ASV105 | 3.62 ± 6.46 | 0.08 ± 0.14 | 3.29 ± 3.55 | 0.07 ± 0.08 | 23.00 ± 25.93 | 0.49 ± 0.56 | - | - |
| | | ASV151 | 15.5 ± 15.46 | 0.33 ± 0.33 | 0.86 ± 2.27 | 0.02 ± 0.05 | 0.00 ± 0.00 | 0.00 ± 0.00 | - | - |
| | | ASV165 | 8.12 ± 6.51 | 0.17 ± 0.14 | 2.00 ± 3.61 | 0.04 ± 0.08 | 0.89 ± 1.27 | 0.02 ± 0.03 | - | - |
| | | ASV178 | 24.00 ± 24.01 | 0.51 ± 0.51 | 1.29 ± 3.40 | 0.03 ± 0.07 | 2.11 ± 5.30 | 0.05 ± 0.11 | - | - |
| | | ASV183 | 0.62 ± 1.77 | 0.01 ± 0.04 | 7.43 ± 6.00 | 0.16 ± 0.13 | 3.89 ± 9.51 | 0.08 ± 0.20 | - | - |
| | | ASV218 | 30.00 ± 47.12 | 0.64 ± 1.01 | 0.14 ± 0.38 | 0.00 ± 0.01 | 0.67 ± 2.00 | 0.01 ± 0.04 | - | - |
| | | ASV233 | 19.62 ± 13.57 | 0.42 ± 0.29 | 0.00 ± 0.00 | 0.00 ± 0.00 | 0.56 ± 1.33 | 0.01 ± 0.03 | - | - |
| | | ASV24 | 103.50 ± 89.08 | 2.22 ± 1.91 | 21.14 ± 34.45 | 0.45 ± 0.74 | 55.67 ± 125.51 | 1.19 ± 2.69 | - | - |
| | | ASV26 | 156.00 ± 108.12 | 3.34 ± 2.32 | 8.00 ± 14.91 | 0.17 ± 0.32 | 59.67 ± 139.21 | 1.28 ± 2.98 | - | - |
| | | ASV269 | 7.25 ± 6.18 | 0.16 ± 0.13 | 1.00 ± 1.41 | 0.02 ± 0.03 | 1.00 ± 2.00 | 0.02 ± 0.04 | - | - |
| | | ASV270 | 11.00 ± 19.50 | 0.24 ± 0.42 | 0.00 ± 0.00 | 0.00 ± 0.00 | 0.11 ± 0.33 | 0.00 ± 0.01 | - | - |
| | | ASV28 | 118.88 ± 53.03 | 2.55 ± 1.14 | 39.43 ± 78.30 | 0.84 ± 1.68 | 17.67 ± 32.20 | 0.38 ± 0.69 | - | - |
| | | ASV285 | 12.12 ± 16.83 | 0.26 ± 0.36 | 0.00 ± 0.00 | 0.00 ± 0.00 | 0.00 ± 0.00 | 0.00 ± 0.00 | - | - |
| | | ASV333 | 11.38 ± 10.53 | 0.24 ± 0.23 | 0.00 ± 0.00 | 0.00 ± 0.00 | 0.33 ± 1.00 | 0.01 ± 0.02 | - | - |
| | | ASV382 | 11.50 ± 13.47 | 0.25 ± 0.29 | 0.00 ± 0.00 | 0.00 ± 0.00 | 0.56 ± 1.67 | 0.01 ± 0.04 | - | - |
| | | ASV472 | 2.38 ± 4.14 | 0.05 ± 0.09 | 0.00 ± 0.00 | 0.00 ± 0.00 | 0.00 ± 0.00 | 0.00 ± 0.00 | - | - |
| | | ASV492 | 5.62 ± 6.25 | 0.12 ± 0.13 | 0.14 ± 0.38 | 0.00 ± 0.01 | 0.78 ± 2.33 | 0.02 ± 0.05 | - | - |
| | | ASV52 | 10.62 ± 13.94 | 0.23 ± 0.30 | 1.14 ± 2.27 | 0.02 ± 0.05 | 0.33 ± 0.71 | 0.01 ± 0.02 | - | - |
| | | ASV53 | 49.88 ± 46.03 | 1.07 ± 0.99 | 4.71 ± 10.81 | 0.10 ± 0.23 | 12.44 ± 22.74 | 0.27 ± 0.49 | - | - |

| | | | | | | | | | | |
|---|---|---|---|---|---|---|---|---|---|---|
| | | ASV557 | 3.25 ± 4.03 | 0.07 ± 0.09 | 0.00 ± 0.00 | 0.00 ± 0.00 | 0.00 ± 0.00 | 0.00 ± 0.00 | - | - |
| | | ASV58 | 49.38 ± 29.22 | 1.06 ± 0.63 | 14.14 ± 25.34 | 0.30 ± 0.54 | 6.11 ± 12.67 | 0.13 ± 0.27 | - | - |
| | | ASV601 | 6.88 ± 7.85 | 0.15 ± 0.17 | 0.00 ± 0.00 | 0.00 ± 0.00 | 0.22 ± 0.67 | 0.00 ± 0.01 | - | - |
| | | ASV607 | 0.00 ± 0.00 | 0.00 ± 0.00 | 0.00 ± 0.00 | 0.00 ± 0.00 | 1.22 ± 1.48 | 0.03 ± 0.03 | - | - |
| | | ASV616 | 5.25 ± 5.18 | 0.11 ± 0.11 | 0.14 ± 0.38 | 0.00 ± 0.01 | 0.00 ± 0.00 | 0.00 ± 0.00 | - | - |
| | | ASV70 | 62.75 ± 59.87 | 1.34 ± 1.28 | 7.57 ± 15.35 | 0.16 ± 0.33 | 2.56 ± 6.58 | 0.05 ± 0.14 | - | - |
| | | ASV74 | 37.12 ± 21.05 | 0.79 ± 0.45 | 13.57 ± 23.40 | 0.29 ± 0.50 | 5.89 ± 12.09 | 0.13 ± 0.26 | - | - |
| | | ASV745 | 3.12 ± 2.64 | 0.07 ± 0.06 | 0.00 ± 0.00 | 0.00 ± 0.00 | 0.11 ± 0.33 | 0.00 ± 0.01 | - | - |
| | | ASV839 | 3.38 ± 4.44 | 0.07 ± 0.10 | 0.00 ± 0.00 | 0.00 ± 0.00 | 0.00 ± 0.00 | 0.00 ± 0.00 | - | - |
| | | ASV99 | 31.12 ± 19.28 | 0.67 ± 0.41 | 10.57 ± 20.83 | 0.23 ± 0.45 | 5.56 ± 12.88 | 0.12 ± 0.28 | - | - |

**Table S11:** Alpha diversity metrics (species richness, diversity, and evenness) for bulk soil (Sb) and rhizosphere (Sr) categorised by treatment (Tx, allium 'A', banana 'B', co-cultivation 'AB' and control 'C'). Significantly different values ($p < 0.05$) are denoted by distinct letters. Species richness is quantified using the Observed, Chao1, and ACE indices, Diversity is assessed via the Shannon and Simpson indices, and Evenness is measured using the Pielou index.

| Gene/ Region | Soil type | Tx | Richness (Observe) | Richness (Chao1) | Richness (ACE) | Diversity (Shannon) | Diversity (Simpson) | Evenness (Pielou) |
|---|---|---|---|---|---|---|---|---|
| 16S | Sb | A | 994.889 ± 209.688 (a) | 1166.653 ± 282.261 (a) | 1174.127 ± 288.960 (a) | 6.127 ± 0.439 (a) | 0.989 ± 0.017 (a) | 0.891 ± 0.054 (a) |
| | | AB | 721.125 ± 199.049 (b) | 766.035 ± 222.735 (b) | 763.436 ± 224.282 (b) | 5.920 ± 0.671 (a) | 0.988 ± 0.025 (a) | 0.903 ± 0.083 (b) |
| | | B | 758.800 ± 194.713 (b) | 812.989 ± 227.816 (b) | 806.129 ± 224.718 (b) | 6.047 ± 0.357 (a) | 0.994 ± 0.006 (a) | 0.917 ± 0.034 (b) |
| | | C | 762.200 ± 196.800 (b) | 809.500 ± 221.991 (b) | 802.632 ± 222.728 (b) | 6.102 ± 0.352 (a) | 0.993 ± 0.010 (a) | 0.924 ± 0.033 (b) |
| | Sr | A | 445.000 ± 41.539 (a) | 745.556 ± 134.761 (a) | 807.618 ± 142.624 (a) | 5.759 ± 0.197 (a) | 0.994 ± 0.004 (a) | 0.945 ± 0.018 (a) |
| | | AB | 404.000 ± 146.632 (a) | 694.595 ± 305.369 (a) | 759.995 ± 352.640 (a) | 5.561 ± 0.691 (a) | 0.992 ± 0.008 (a) | 0.946 ± 0.016 (a) |
| | | B | 456.714 ± 28.768 (a) | 754.790 ± 75.773 (a) | 824.071 ± 77.216 (a) | 5.829 ± 0.091 (a) | 0.995 ± 0.001 (a) | 0.952 ± 0.007 (a) |
| ITS1 | Sb | A | 47.250 ± 9.270 (a) | 81.908 ± 24.581 (a) | 91.952 ± 18.303 (a) | 3.154 ± 0.518 (a) | 0.889 ± 0.074 (a) | 0.818 ± 0.093 (abc) |
| | | AB | 46.100 ± 17.929 (a) | 72.115 ± 36.440 (a) | 76.166 ± 37.794 (a) | 3.255 ± 0.761 (a) | 0.914 ± 0.085 (a) | 0.881 ± 0.078 (a) |
| | | B | 40.400 ± 10.069 (a) | 88.196 ± 33.481 (a) | 85.482 ± 17.983 (a) | 2.862 ± 0.646 (a) | 0.856 ± 0.116 (a) | 0.772 ± 0.126 (bc) |
| | | C | 35.100 ± 16.736 (a) | 76.852 ± 44.519 (a) | 88.544 ± 42.515 (a) | 2.429 ± 1.083 (a) | 0.744 ± 0.282 (a) | 0.675 ± 0.234 (bc) |
| | Sr | A | 131.556 ± 48.603 (a) | 143.785 ± 53.007 (a) | 144.482 ± 53.134 (a) | 3.359 ± 1.144 (a) | 0.853 ± 0.201 (a) | 0.695 ± 0.175 (a) |
| | | AB | 103.667 ± 47.513 (b) | 121.631 ± 54.293 (a) | 123.288 ± 54.944 (a) | 2.448 ± 1.285 (ab) | 0.684 ± 0.310 (ab) | 0.517 ± 0.241 (ab) |
| | | B | 96.000 ± 29.394 (ab) | 132.387 ± 51.660 (a) | 135.174 ± 49.345 (a) | 2.040 ± 0.516 (b) | 0.675 ± 0.143 (b) | 0.448 ± 0.089 (b) |
| ITS2 | Sb | A | 165.000 ± 44.475 (a) | 189.641 ± 57.945 (a) | 195.921 ± 63.551 (a) | 3.702 ± 0.691 (abc) | 0.899 ± 0.092 (abc) | 0.729 ± 0.114 (ab) |
| | | AB | 161.889 ± 34.090 (a) | 175.808 ± 39.692 (a) | 180.457 ± 41.888 (a) | 3.944 ± 0.371 (a) | 0.946 ± 0.032 (a) | 0.779 ± 0.068 (a) |
| | | B | 131.125 ± 34.498 (a) | 174.722 ± 49.506 (a) | 182.764 ± 55.632 (a) | 3.006 ± 0.765 (bc) | 0.840 ± 0.112 (bc) | 0.619 ± 0.137 (bc) |
| | | C | 136.444 ± 47.584 (a) | 170.114 ± 57.689 (a) | 173.069 ± 54.570 (a) | 3.014 ± 0.656 (bc) | 0.846 ± 0.074 (bc) | 0.616 ± 0.089 (c) |
| | Sr | A | 148.125 ± 27.126 (a) | 154.841 ± 29.257 (a) | 154.652 ± 29.394 (a) | 3.815 ± 0.529 (a) | 0.942 ± 0.042 (a) | 0.766 ± 0.097 (a) |
| | | AB | 110.714 ± 45.792 (a) | 129.645 ± 41.810 (a) | 127.653 ± 41.236 (a) | 2.553 ± 0.965 (b) | 0.767 ± 0.195 (b) | 0.540 ± 0.165 (b) |
| | | B | 114.000 ± 53.486 (a) | 132.723 ± 53.249 (a) | 138.142 ± 52.254 (a) | 2.536 ± 0.988 (b) | 0.766 ± 0.162 (b) | 0.536 ± 0.160 (b) |
